# A protein language model links the *E. coli* pangenome functional landscape geometry to host proteostasis

**DOI:** 10.64898/2026.01.15.699719

**Authors:** Daniel Martinez-Martinez, Cassandra Backes, Aleksandra Zečić, Andreea Aprodu, Hannah M. Doherty, Jonas Widder, Andrew Ivan, Laurence Game, Iliyana Kaneva, Georgia Roumellioti, Alex Montoya, Holger Kramer, Filipe Cabreiro

## Abstract

Understanding how bacterial diversity at strain level resolution shapes host physiology is a central challenge in microbiome research. The vast, functionally unknown genetic diversity within a species pangenome makes it difficult to connect genes to function and their impact on host physiology. Here, we explore how the functional landscape of the *Escherichia coli* pangenome impacts transcriptional responses in *Caenorhabditis elegans* and show that traditional gene-centric methods fail to provide significant functional associations with the host. Thus, we developed a pangenome framework that leverages the protein language model ProtT5 and generates unique strain embeddings representing the functional potential of each 9,558 *E. coli* isolate. Stratification of the pangenome into distinct functional guilds aligned with key host processes such as cell division, metabolism and proteostasis. Through mGWAS analysis we identify a critical interplay between the extensive network of bacterial chaperones and proteases in regulating host proteostasis. We find that the bacterial chaperone DNAK/HSP70 and protease ClpX fine-tune the host ubiquitin-proteasome system by controlling propionate and vitamin B12 availability. These findings reveal a conserved ‘co-proteostasis’ mechanism as a key phenomenon modulating host-microbe interactions through metabolic communication. Our pangenome-to-phenotype approach offers a powerful strategy to decode bacterial pangenome functional diversity, directly linking microbial genomic variation to host proteostasis.

## INTRODUCTION

Protein aggregation is a defining feature of multiple age-related neurodegenerative disorders and a hallmark of ageing ^1^. Misfolded protein accumulation and aggregate formation signify a failure of the proteostasis network that normally safeguards protein folding, trafficking, and degradation, with a key role for the ubiquitin–proteasome system (UPS), a central pathway for selective clearance of damaged or misfolded proteins ^2^. Although cell-intrinsic proteostasis pathways, including the ubiquitin–proteasome system, have been dissected in considerable detail^3^, it remains unclear how systemic cues, particularly those originating from host–microbe interactions, shape organismal proteostasis capacity.

The gut microbiome is emerging as a central modulator of host metabolism with microbes exerting broad control over metabolite systemic abundance and distribution. For example, microbiota-derived metabolites can reprogram metabolic homeostasis, mitochondrial activity, oxidative stress resilience, and proteostasis pathways ^4–7^. Thus, despite the metabolic capacity of the host being vastly expanded by its resident microbiome, correlating specific microbial signatures with physiological outcomes remains a fundamental challenge. Animal models such as *Caenorhabditis elegans* have been successfully repurposed as biosensors to study the mechanisms underlying host-microbe interactions^8–10^. Indeed, recent studies demonstrate that specific microbial taxa can attenuate protein aggregation in organismal models. In *Caenorhabditis elegans*, *Bacillus subtilis* suppressed α-synuclein aggregation linked to Parkinson’s disease, whereas the absence of curli proteins in *Escherichia coli* reduced polyglutamine aggregation associated with Huntington’s disease ^11,12^. However, while strain-specific impacts on host physiology are increasingly recognized, the vast genetic diversity within individual bacterial species remains largely underexplored. Current strategies relying on phylogenetic markers or linear reference genomes fail to fully capture this functional potential, leaving a gap in our ability to predict how intra-species genetic heterogeneity drives distinct host phenotypes^13^. The *E. coli* pangenome represents a vast reservoir of uncharted metabolic potential given its ecological ubiquity and open genomic architecture^14,15^. For instance, distinct *E. coli* strains elicit divergent host responses through differential production of metabolites, such as vitamin B12 or betaine^16–18^. Yet, standard laboratory strains, often used to study these interactions capture only a minute fraction of this natural diversity^19^. As a result, new analytical frameworks are needed that move beyond sequence identity to capture the latent functional potential of bacterial proteomes and map them directly to host physiology, thereby bridging the gap between reductionist models and the complex reality of natural microbiomes^20^.

Here, we bridge this gap by combining high-throughput transcriptomics of *C. elegans* with a novel machine-learning approach that utilizes the protein language model (pLM) ProtT5^21^ to recreate the functional landscape of a pangenome of 9,558 *E. coli* assemblies. Protein embeddings in the latent space from the ProtT5 model have shown to be accurate and informative representations ^22^, a requirement given the uncharted genomic content of the *E. coli* pangenome. We integrated the geometrical representation from the pLM with the genetic background of the strains to generate strain embeddings, vector representations encapsulating the total functional potential of a bacterial strain. By exposing *C. elegans* to a diverse library of 592 *E. coli* strains, we demonstrate that the geometry of the bacterial embedding space accurately predicts host phenotypic variance, revealing a profound link between the microbial pangenome, microbial proteases and host proteostasis. We identify a cross-domain co-proteostasis mechanism where the bacterial proteostasis network (DnaK/ClpX) regulates vitamin B12 and propionate metabolism, dictating metabolic rewiring in the host through B12-dependent or independent metabolic shunt that regulates host ubiquitin-proteasome system function.

## RESULTS

### The *E. coli* pangenome elicits a vast range of transcriptomic profiles in *C. elegans*

*E. coli* is known to have an open pangenome^15^, meaning that the various strains within the *E. coli* species contain unique genes coding for proteins whose functions are essential for their distinct functional properties. Thus, we hypothesized that this extensive genetic variation within the *E. coli* pangenome would translate to defined functional properties dictating host responses. To interrogate these host responses to individual bacterial strains, we generated high-resolution bulk RNAseq transcriptional profiles for 592 distinct *E. coli* – *C. elegans* mono-association pairs (Figure 1A, Table S1). We curated a library combining the EcoRef collection and additional strains with broad phylogenetic coverage (Figure 1B)^15,19^. This panel spans the major *E. coli* phylogroups - evolutionary lineages defined by specific gene markers that are traditionally linked to distinct ecological roles, and primarily comprises commensal strains isolated from human and animal hosts (Figure 1C; Figure S1A), with roughly 50% belonging to phylogroup B2. Analysis of the strain genomes confirmed an open pangenome architecture: a conserved core of 3,265 gene families (>95% presence), a shell genome of 2,589 gene families (15% - 95% presence), and a diverse and large cloud genome of 20,113 rare gene families (<15% presence) (Figure 1D). This distribution highlights the immense reservoir of genetic diversity available to influence host physiology. Next, we profiled the host response by raising synchronized *C. elegans* (N2) on each bacterial strain and sequencing total RNA from Day 1 adults. Following rigorous quality filtering and batch effect correction (Figure S1B-D), we established a robust transcriptional dataset comprising 16,410 unique genes. This yielded high-coverage data with an average of 8,545 ± 254 transcripts detected per sample (Figure S1E). Remarkably, we found that the commonly used laboratory strain *E. coli* OP50 used for most studies in classic genetics and aging-related publications induces a transcriptional profile in *C. elegans* distinct to the transcriptional signatures of the majority of strains (Figure 1E), including the K-12 MG1665 lab strain whose genome was one of the first *E. coli* reference sequences to be completed and extensively curated.

**Figure 1.**
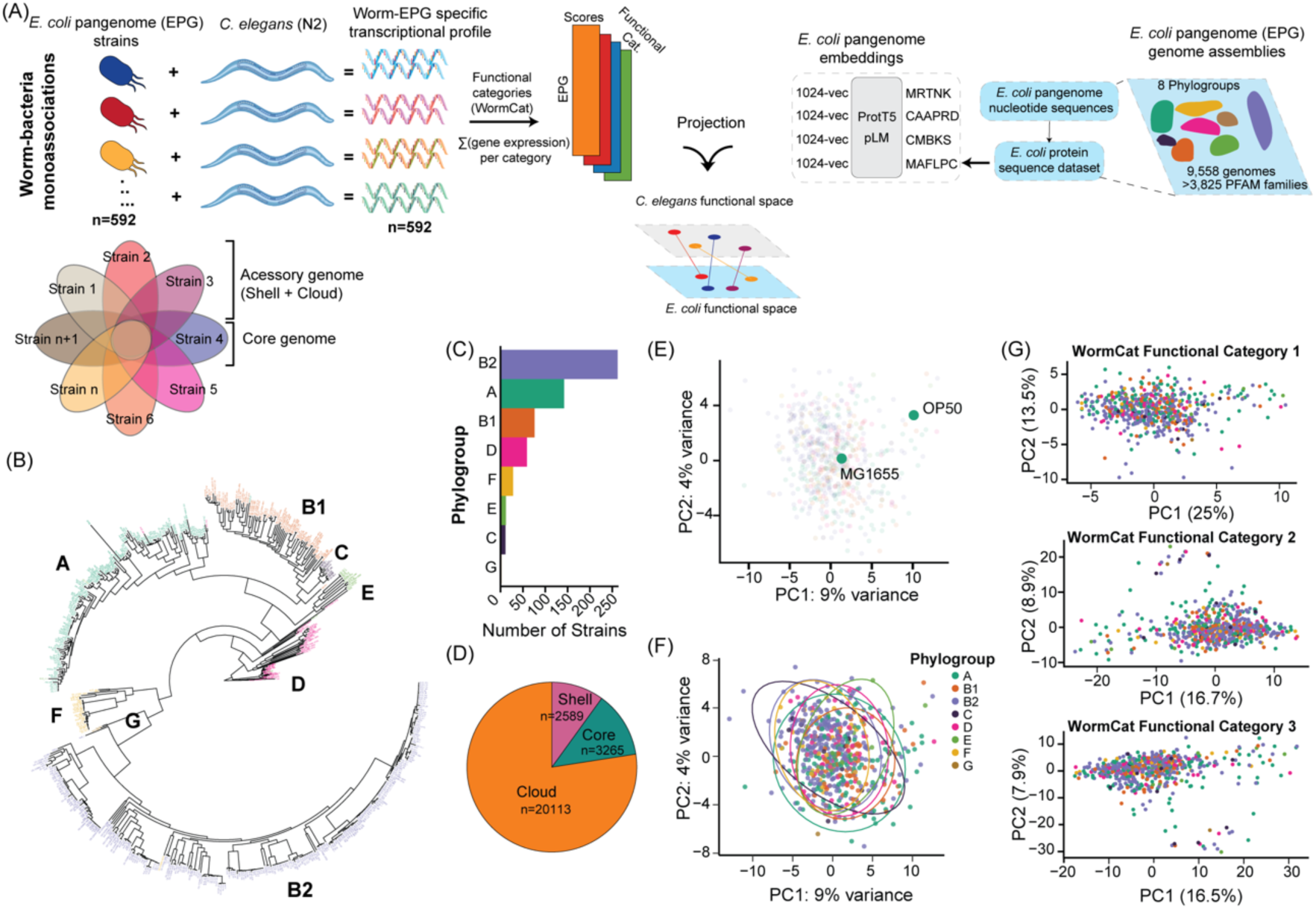
The physiology of *C. elegans* is regulated at the *E. coli* pangenome scale. (A), Schematic representation of experimental and analytical workflow: *C. elegans* transcriptomes were profiled for each of the 592 *E. coli* strains and then summarized in WormCat functional categories. The *E. coli* pangenome was calculated for 9,558 strains and its linear reference was used to geometrically represent the functional potential per strain with the pLM ProtT5. Both biological layers were used to map host-microbe functional landscapes. (B), Phylogenetic tree computed from the core genome fraction of the 592 *E. coli* strains panel with tips colored following the major phylogroups. The tree branch length reflects genetic distances. (C), Pie chart showing the distribution of gene families belonging to the core (>95% presence), shell (95% to 15% presence), or cloud (<15% presence) genome across the 592 *E. coli* strains. (D), Distribution of *E. coli* phylogroups across the 592 *E. coli* strains. (E), Representation of the transcriptional distance between two common *E. coli* lab strains belonging to the same phylogroup, MG1655 and OP50, which are known to induce differences in the worm physiology. (F), Principal component analysis (PCA) of the whole *C. elegans* transcriptional profiles. Each point representing animals raised on a single *E. coli* strain and colored by the phylogroup of the corresponding strain. (G), PCA of WormCat level 1, level 2 and level 3 categories, depicting the functional landscape of *C. elegans* transcriptional responses to the *E. coli* strain panel. Each point represents the WormCat functional category for a given strain and colored by the phylogroup of the corresponding *E. coli* strain.

Next, we investigated whether grouping strains by phylogroup, as a proxy for bacterial function, would reveal a structure in the worm transcriptional response to the *E. coli* strain panel. The whole transcriptional profiles were correlated to the phylogroup partitioning of the *E. coli* pangenome and variance explained by this functional division was measured. Principal component analysis (PCA) revealed a modest separation between the phylogroups included in this screening (Figure 1F), consistent with a weak clustering of pairwise Euclidean distances between strains (Figure S1F). Moreover, a permutational analysis of variance (PERMANOVA) for the full transcriptome dataset indicated a significant effect of phylogroup. However, the model explained only approximately 1.5% variance. Nevertheless, the large Euclidean distance in transcriptional responses observed between strains known to elicit a distinct physiological response in the worm such as OP50 and MG1655 (Figure 1E), suggests that a robust biological signal exists within this transcriptional landscape. To facilitate mapping the worm response onto the *E. coli* pangenome, we reasoned that clustering the normalized read counts to discrete functional categories would improve our ability to map worm response. For this, the curated worm database from Holdorf *et al*.^23^ was leveraged and normalized read counts were aggregated for all genes within each functional category at the three defined hierarchical levels defined in the database (Figure 1A, Table S2). This yielded three matrices of increased granularity, ranging from 33 broad categories (level 1) to 461 highly specific functional categories (level 3). This stratification generated a dense phenotypic landscape comprising 272,912 phenotypic worm data points at level 3 functional category resolution. PCA for each category (Figure 1G) revealed a complex landscape where simple patterns remain elusive, underscoring the difficulty of mapping pangenome structure onto worm responses using traditional means. While PERMANOVA testing for level 3 functional categories identified a statistically significant effect (P= 0.041), the low variance explained (1.45%) highlights that evolutionary history alone is a minor driver of host phenotype. Together, these results show that *C. elegans* mounts highly strain-specific responses to *E. coli* strains, both at the whole transcriptional level and at the functional category levels. This confirms that the openness and complexity of the *E. coli* pangenome pose a high-dimensional biological challenge that cannot be fully captured by a standard phylogenetics approach.

### The functional content of *E. coli* is encoded in the protein embedding space

The genetic repertoire of commonly used laboratory strains represents only a fraction of the total species diversity ^14^. However, there is a consensus supporting that *E. coli* clades, whether classified from multi-locus sequence typing (MLST) or broader phylogroup classification, harbor characteristic functional enrichments associated with their primary ecological niche^15,19,24^. Given the shortcomings of current approaches, to systematically investigate the functional diversity of the *E. coli* pangenome, we established a comprehensive panel of genome assemblies to fully capture this functional richness. We accessed the NCBI Genomes database (Jan 2024) and selected 8,829 high-quality genome assemblies, to which we added assemblies for the strains available in our laboratory, yielding a total of 9,558 high-quality genomes (see Methods, Figure S1G). Phylogroup proportions in this extended collection followed a similar pattern as in our laboratory panel (Figure 2B, R = 0.82, p = 0.014, Figure S2A). This resulted in a total of 92,435 gene families split into a core of 3,005 genes, a shell of 2,917 genes, and a cloud of 86,513 genes (Figure 2A, Table S3). Consistent with previous work, the *E. coli* pangenome remained open as indicated by the Heap’s law fit (g = 0.28, Figure S2B). However, we observe that new functions start to saturate, with the core genome rapidly stabilizing. Additionally, the pairwise genetic similarity for this larger panel is similar to our laboratory panel (median Jaccard similarity of 0.641 and 0.65 respectively, Figure S2C). Phylogeny on the core genome shows that *E. coli* robustly follows the phylogroup evolutionary lineages (Figure 2C). When considering gene presence-absence patterns within the cloud genome as a proxy for differential functional content, strain relationships strongly follow the same phylogenetic structure (Figure 2D). This congruence between core genome phylogeny and total gene content demonstrates that a shared evolutionary history shapes the full genomic repertoire of *E. coli*, linking deep ancestry to functional gene repertoire at the strain level.

**Figure 2.**
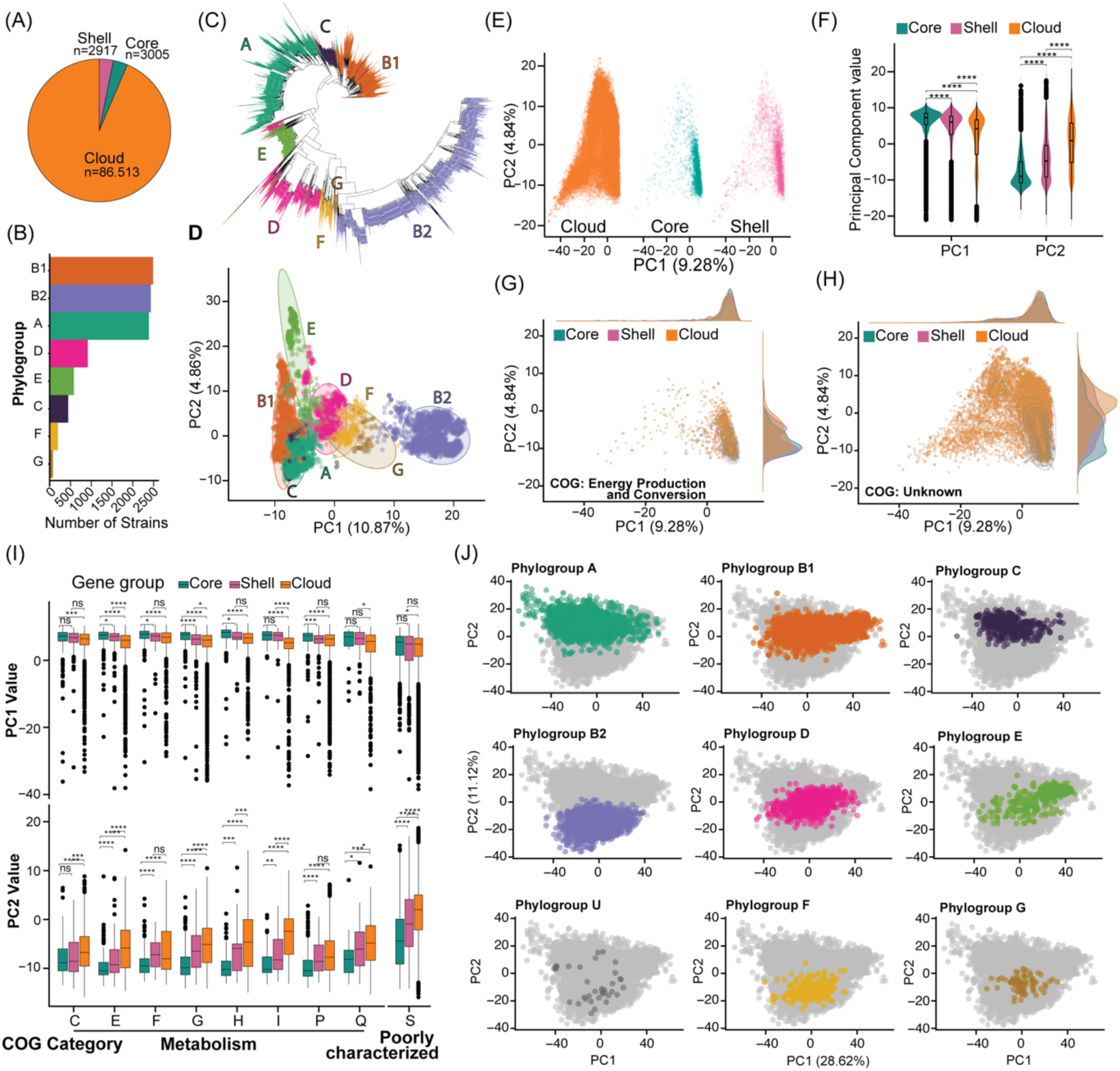
The functional landscape encoded in the *E. coli* pangenome can be leveraged to create a functional map of the species. (A), Pie chart showing the number of gene families assigned to the core (>95% presence), shell (95% to 15% presence), or cloud (<15% presence) genome across the 9 558 *E. coli* strains. (B), Distribution of *E. coli* phylogroups across the 9 558 *E. coli* strains. (C), Phylogenetic tree computed from a set of 275 conserved genes from the core genome of the 9,558 *E. coli* strains. Tips are colored following the major phylogroups and the tree branch lengths reflect genetic distances. (D), PCA of gene presence/absence across the cloud genome of the *E. coli* strain panel. Each point represents a strain, colored by phylogroup. (E), PCA projection of protein embeddings with Orange/Pink/Green representing genes belonging to the cloud/shell/core genome respectively. (F), Violin plots with box plots embedded representing the distribution from the genome fractions per principal component (n = 92,244; p<0.001, One-way ANOVA). (G), PCA projection of protein embeddings for genes belonging to the COG category C, Energy Production and Conversion. (H), PCA projection of protein embeddings for genes belonging to the COG category S, Unknown. (I), Box plots of the PCA coordinates per COG category belonging to the major category of Metabolism and Poorly Characterized (n = 33-18,050). (NS P > 0.05, *P < 0.05, **P < 0.01, ***P < 0.001, two-tailed pairwise T-test with BH). (J), PCA projection of the 9,558 *E. coli* strain embeddings, colored and split by their respective phylogroups.

To characterize functional content, four gene ontology (GO) annotation strategies were benchmarked by leveraging the linear reference of representative gene families. The most widely used methods rely on sequence similarity which favors accuracy at the cost of limited range. Recently, the development of methods relying on machine learning (ML) models have been proven to achieve significant improvements in quality and coverage^25^. The methods to predict GO terms included Interproscan^26^ and eggNOG-mapper^27^, which are based on sequence similarity, and ML methods such as Proteinfer^28^, a Convolutional Neural Network method, and GoPredSim^29^, which leverages the protein language model (pLM) ProtT5-XL-U50^21^ by creating an embedding representation for each protein. ML-based approaches annotated substantially more genes per genome than sequence-similarity based methods (41.7-52.7% increase, Figure S2D, p<0.001, one-way ANOVA). This difference was primarily driven by annotations in the cloud and shell genomes where unique sequences are mostly found (Figure S2E,F). Despite this significant improvement, 2,873 gene families remained unannotated. Exploiting the hierarchical information contained in GO annotations, an analysis of the maximum information content per gene and method revealed that the GoPredSim, which leverages the pLM ProtT5 embeddings, produced the most informative annotations (Figure S2G). These results underscore that much of the accessory functional landscape from *E. coli* remains largely uncharacterized but can be more effectively represented using pLM embedding-based models, given the prediction abilities and breadth of information.

Protein embeddings from the ProtT5-XL-U50 pLM were then used to create a comprehensive functional map of the *E. coli* pangenome (Figure 1A, Figure S1G). PCA of the resulting high-dimensional embeddings revealed a spatial organization aligned with pangenome structure (Figure 2E, Figure S2H). The core genome occupies a compact, dense region of the space, which expands through the shell genome and into the vast, sparsely populated region occupied by the cloud genome with significant differences between these compartments (Figure 2F, Figure S2H, p < 0.001, one-way ANOVA). This geometric arrangement demonstrates that the embedding space quantitatively captures functional diversity, transitioning from the conserved core functions to the diverse and mobile cloud genome. Analysis of the COG categories further show that core-associated functions such as energy production and conversion (C) or cell cycle control (D) are enriched in the dense core region. However, functions known to be environment-dependent such as carbohydrate transport (G), defense mechanisms (V) or poorly characterized and unknown (S) are shifted towards the periphery (Figure 2G,H,I, Figure S 2I,J,K). Collectively, these results establish that the geometry of the protein embedding space created through the pLM ProtT5 acts as a quantitative proxy for the pangenome’s functional architecture.

Given that the functional content of a pangenome is encoded in the geometry of the protein embedding space, we extended this framework to establish a microbial strain identity, defined by the combination of its conserved core functions and unique set of accessory functions. Each strain was embedded as a single vector by combining binary gene content (presence/absence) with the corresponding protein embeddings, calculated as an average of all protein vectors considering the whole gene content (see Methods section), yielding a unique representation per strain that determines their functional potential. PCA of these strain embeddings produced a pangenome-level functional landscape in which strains are positioned according to functional capacity (Figure 2J, Figure S2L, Table S4). The principal component analysis of the strain embeddings revealed a structured functional landscape that validates our model while highlighting the limitations of pure phylogeny approaches (p < 0.001, one-way ANOVA, Figure S2L). PC2 (11.12%) cleanly separated phylogroups, confirming that the embeddings correctly encoded the strains’ evolutionary history, whereas PC1 (28.62%) was driven by within-phylogroup variability. This analysis proves that the vast majority of *E. coli* functional diversity is driven by strain-specific adaptations. This pattern is captured by alternative strain embedding pooling methods and representations, supporting the robustness of these findings (Figure S3A-D). We observe the strain embeddings recapture the clusters defined by ecology and evolution where phylogroups A, B1 and C formed a clearly defined cluster, followed by a distinct cluster composed of phylogroups D, E, F and G bridging towards the phylogroup B2, which occupied a distinct, distant region. This separation of phylogroups along PC2 supports the evolutionary history between the phylogroups (Figure 2J). Together, these results demonstrate that protein language model embeddings provide a powerful, compact representation of both genes and whole genomes, enabling the encoding of *E. coli* pangenome structure, the delineation of conserved versus accessory functions, and the representation of entire strains as single, functionally meaningful vectors that can be directly linked to host phenotypes.

### Host physiology maps onto the strain embeddings functional landscape

We next hypothesized whether using the *E. coli* strain embeddings would improve our ability to establish causal links between microbial functions and host physiology at the pathway level. To this end, we leveraged the geometry of the PCA projection coordinates derived from the strain embeddings as a functional map to position *C. elegans* transcriptional and functional categories (Figure 1A). The strain embedding map for the subset of *E. coli* strains used in mono-association experiments recapitulated the distinctive structure observed for the full pangenome map, revealing a clear separation within and between phylogroups along PC1 and PC2 coordinates, respectively (Figure 3A). To connect the *E. coli* pangenome functional landscape and worm physiology, we calculated the Spearman correlations between the level 3 pathway categories in the worm transcriptome to the PC1 and PC2 coordinates of the strain embeddings. This analysis revealed a distinct set of host functional categories regulated at the pangenome level (Figure 3B). Notably, all significant correlations between host pathways and strain embedding axes were linked to PC2, which separates the distinct phylogroups in the pangenome (Figure 3C, Table S5). The significant pathways involved in the host physiology regulation at the pangenome level, clustered into biologically broad processes including cell cycle, central and one-carbon metabolism, proteostasis and response to stress (Figure 3D, Figure S3E, p < 0.05, BH correction). The prediction of worm phenotype at level 3 pathway categories from the strain embeddings using Random Forest yielded a similar outcome, emphasizing the robustness of the methodology and the biological signal obtained (Figure S3F). These processes have been formerly linked to the regulation of several host phenotypes, including development and aging^30–32^. Given the *E. coli* pangenome functional landscape - host physiology connections, we sought to validate these associations experimentally. We performed a reporter-based quantitative screen of 13 fluorescent transcriptional or translational reporters representing key genes within the main pathways identified in the functional mapping (Figure 3E,F, Table S6, Table S11). We confirmed the robustness of the experimental pipeline by testing two independent biological replicates over 589 *E. coli* pangenome strains for each reporter (Pearson correlation > 0.7, Figure S3G). We then mapped the reporter gene expression levels to the *E. coli* strain embeddings and found that all 13 gene reporter expression profiles significantly correlated with the strain embedding geometrical projection, mirroring the transcriptional landscape pattern observed (Figure 3G, p < 0.05, BH correction). Together, these results show that the *E. coli* functional landscape can be quantitatively mapped onto the host transcriptional and physiological programs by leveraging strain embeddings, uncovering bacterial-driven regulation of fundamental host processes such as central and 1CC metabolism, stress response, or proteostasis.

**Figure 3.**
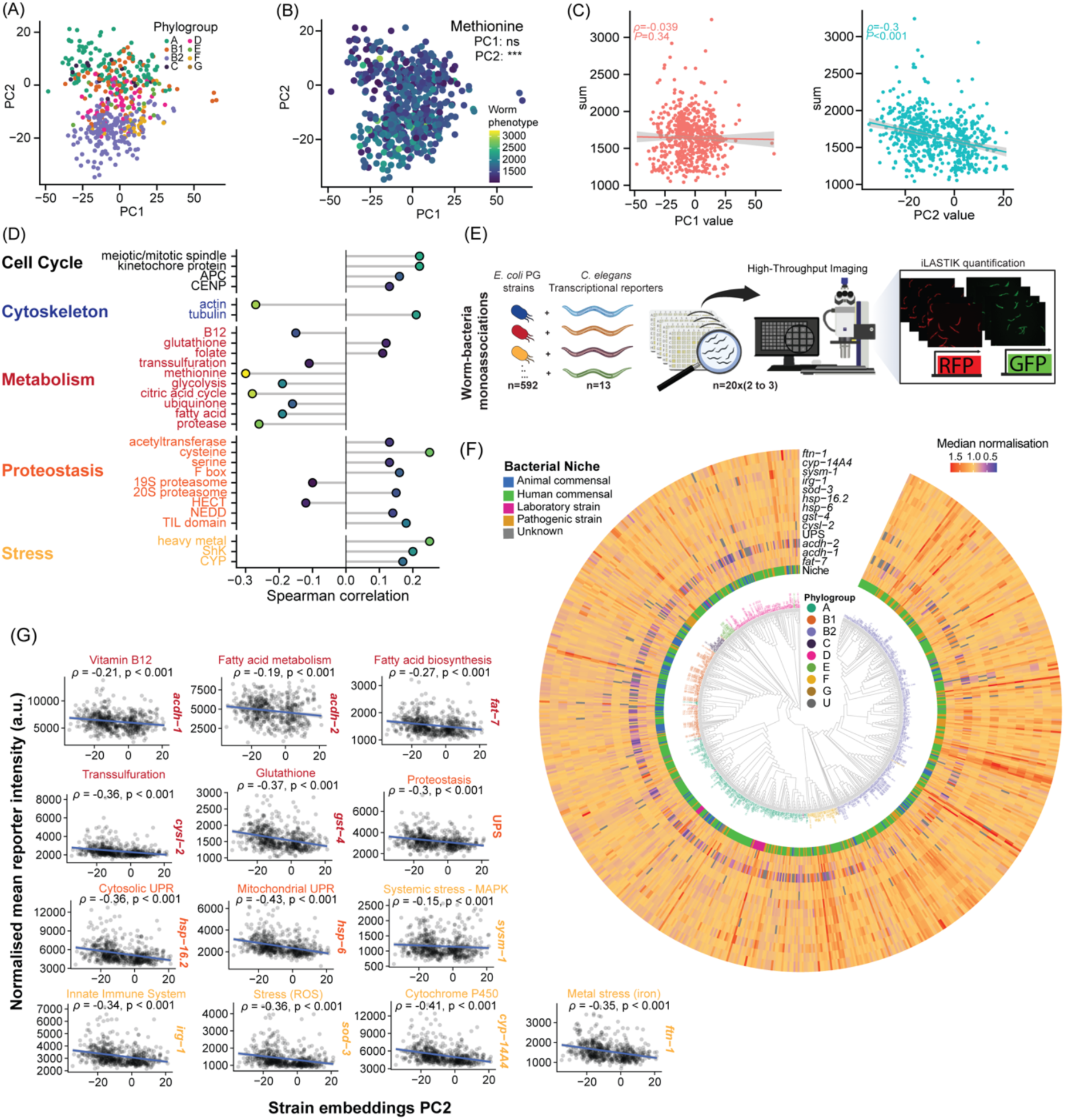
*E. coli* strain embeddings map bacterial functional landscape onto host functional responses. (A), PCA projection for the 592 *E. coli* strain embeddings included in the RNA-seq screen with points colored by phylogroup. (B), PCA projection of the 592 *E. coli* strain embeddings with dots colored by Methionine functional scores derived from *C. elegans* transcriptomes. (n = 592, Spearman correlation, *P < 0.05, **P < 0.01, ***P < 0.001). (C), Spearman correlation plots for the two Principal Components with the Methionine functional scores. Spearman correlation coefficient is represented as the ρ value (n = 592). (D), Bubble plot summarizing Spearman correlation coefficients (ρ) for significant WormCat functional categories and PC2 of the strain-embedding PCA across the 592 strains. (E) Diagram showing the high-throughput experimental design to analyze *E. coli* pangenome monoassociations with *C. elegans* transcriptional gene reporters. (F) Phylogenetic tree of the *E. coli* strain pangenome panel annotated with ecological niche and *C. elegans* fluorescent reporter responses. The innermost layer indicates the origin of each strain (human commensal, human pathogenic, animal commensal, laboratory strain or unknown). Outer layers show median normalized fluorescence ratios for each *C. elegans* gene reporter, mapped onto the corresponding *E. coli* strain. (G), Spearman correlations between the PC2 of the strain embeddings PCA and biologically relevant *C. elegans* gene reporter phenotypes. Each facet shows the relationship between PC2 of the *E. coli* strain-embedding PCA (x-axis) and normalized mean reporter intensity from the high-throughput imaging screen (y-axis) for each *C. elegans* gene reporter. Spearman correlation coefficients (ρ) and significance are represented for each case.

### Microbial GWAS analysis links bacterial proteostasis to host proteostasis

Next, we leveraged these data to obtain further insights between microbe-host functional relationships. Plotting reporter expression by strain revealed distinct transcriptional programs which were largely independent of their phylogenetic relatedness (Figure S4A). Similarly, pairwise strain correlations of reporter activity indicated that bacterial functional guilds transcended phylogenetic relatedness on regulating specific host transcriptional responses (Figure 4A,B, Figure S4A). Interestingly, we observed a global bias towards positive associations between strain pairs (∼65%, Figure S4B,C), implying a shared host transcriptional response across the *E. coli* pangenome. Consistent with this, we observed significant pairwise correlations between the 13 reporters (Figure 4C). Together, these findings suggest that functional variation across the *E. coli* pangenome encodes regulatory signals that can modulate core host physiological programs. Given that the worm processes found in the pangenome functional mapping act concomitantly influencing higher processes like ageing, we next wondered whether there was a shared bacterial genetic program affecting the worm pathways at the pangenome level. To identify these bacterial pangenome features, we employed microbial genome wide association studies (mGWAS) analytical approaches to map the genetic variation from the pangenome at unitigs level. By focusing the mGWAS analysis to the significant worm pathways found from the mappings of the strain embeddings functional landscape (Figure 3D), we obtained lists of significant *E. coli* gene markers per worm category. A permutational analysis of the Jaccard similarity of the pangenome gene markers per worm category highlighted a strong interconnectedness between the markers affecting the worm proteostasis and one-carbon metabolism processes as a shared bacterial driver for host physiology (Figure 4D, Figure S4D, p < 0.05, BH correction). KEGG functional enrichment analysis on statistically significant *E coli* genes associated with changes in host proteostasis and proteolysis functions together with one-carbon metabolism (Figure 4D, S4D), highlight lipopolysaccharide biosynthesis and flagellar assembly processes or O-Antigen nucleotide sugar biosynthesis/colanic acid metabolic processes where unitigs in genes belonging to these functional categories regulate host proteostasis (Figure S4E). Further, the network representing protein interactions from the significant genes analyzed shows that the bacterial protease ClpX has the highest average effect size suggesting that unitigs from this gene are associated with higher phenotypic values (Figure 4E, Table S7). Since colanic acid production is also regulated by the bacterial protease Lon to regulate *C. elegans* mitochondrial unfolded protein response^33,34^, we hypothesized that bacterial proteostasis comprised of proteases and chaperones may regulate host proteostasis.

**Figure 4.**
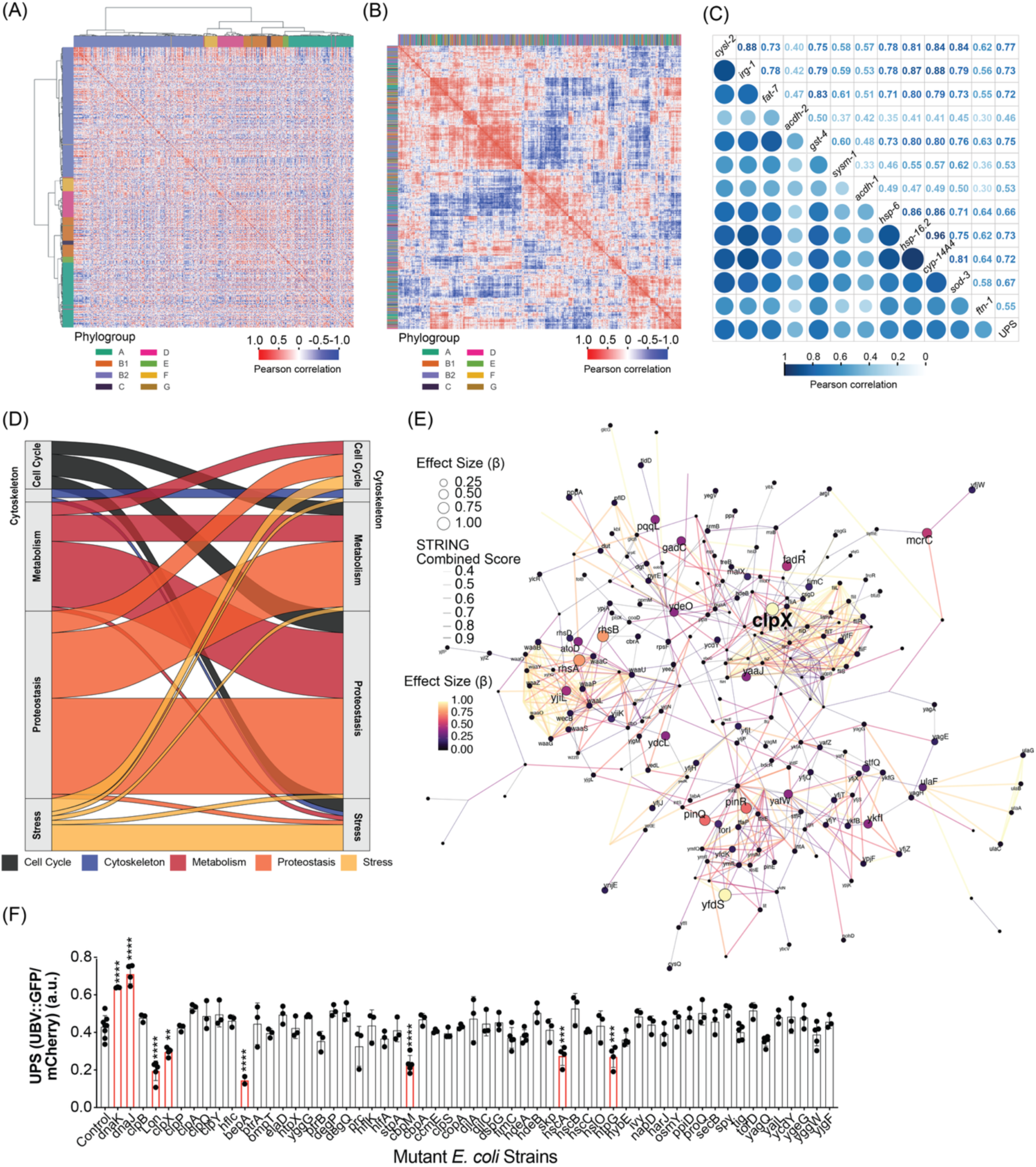
Bacterial strains possess a shared genetic program affecting host proteostasis. (A, B), Strain-strain Pearson correlation heat map derived from *C. elegans* transcriptional gene reporters. Strains are clustered by their phylogenetic distances (A) or by their host fluorescent reporter activity (B) and colored by phylogroup (n = 589). (C), Pearson correlation between *C. elegans* transcriptional gene reporter data shown as circles (lower triangle) and correlation values (upper triangle) (n = 589). (D), Sankey diagram representing the interconnection of worm processes (color-coded) affected by shared bacterial genetic drivers calculated from permutational Jaccard distances from the mGWAS analysis. (E) Protein-protein interaction network obtained from the genes captured in the shared response from the mGWAS analysis. The network topology represents the interactions from STRING DB based on the *E. coli* K-12 model, with edge color and size representing the STRING combined score, and node size and color representing the squared effect size (β) captured by mGWAS. Labels from nodes with β >0.25 are shown. (F) Normalized brightness of the worm reporter Ub(G76V)–GFP as a ratio of GFP over mCherry (UPS fluorescence) fed on *E. coli* knock-out strains for proteins involved in bacterial proteostasis (n=3-8). Stats correspond to comparison against the control (one-way ANOVA) represented as *P < 0.05, **P < 0.01, ***P < 0.001, ****P < 0.0001, NS P > 0.05.

In eukaryotic cells, the ubiquitin-proteasome system (UPS) plays a central role in maintaining proteostasis by controlling the degradation of damaged proteins. In *C. elegans* the functional state of UPS is assessed using the UPS reporter strain expressing the ubiquitously driven ubiquitin-fusion degradation substrate Ub(G76V)–GFP, whose accumulation reflects impaired ubiquitin–proteasome-system–mediated protein turnover^35^. Host UPS displayed strong bacterial strain-dependent variation, as observed for the *E. coli* pangenome (Figure 3F-G) and *E. coli* laboratory strains^35^, as well as on individual bacterial isolates from distinct phyla of the *C. elegans* microbiome (Figure S4F, Table S8)^36,37^, suggesting fine levels of mechanistic regulation. Given our pLM and mGWAS combined analysis pointing to bacterial proteases regulating the host UPS, we quantitatively tested all known *E. coli* proteases and chaperones in their ability to regulate host UPS (Figure 4F). We confirmed that deletion of Lon, ClpX significantly reduced UPS fluorescence as well as the BepA protease and HtpG, HscA, CbpM chaperones (Figure 4F). Surprisingly, deletion of the functional DnaK and DnaJ heat shock pair increased UPS fluorescence. Overall, these data suggest that host UPS is regulated by the function of both bacterial proteases and chaperones genes widely conserved in the *E. coli* pangenome.

### Propionate and vitamin B12 at the interface of bacterial-host “co-proteostasis”

First, we confirmed our screen findings with measurements at the protein level and found a robust and significant response for the proteases and chaperones identified in our screen (Figure 5A, Figure S5A-C, Table S9). To better understand the regulatory networks between these chaperones and proteases and the potential mechanisms involved in the regulation of host proteostasis, we performed proteomics of each individual mutant strain. Deletion of *dnaK* led to pronounced changes in the proteome landscape followed by the deletion of *lon* and *clpX* (Figure S5D). We observed an intricate compensatory mechanism whereby the single deletion of any of these proteases or chaperones leads to significant changes in a network of other chaperones and proteases (Figure 5B, Figure S5E, Table S10), with greater effects observed for *dnaK* and *clpX* mutants. Thus, we tested whether co-regulatory effects of proteases and chaperones could regulate host UPS response through the creation of double-mutant *E. coli* strains of *dnaK*. Notably, only the combined loss of *dnaK* and *clpX* abolished the UPS activation induced by the *dnaK* mutant alone (Figure 5C, Figure S5F,G) without compromising bacterial fitness (Figure S6A). U.V-irradiation experiments to abolish metabolic activity of *E. coli* and alter their metabolome^38^, further show that UPS regulation was dependent on the active metabolism of *E. coli* (Figure 5D). To identify the potential mechanism(s) responsible, we performed proteomics in both single and double *E. coli* mutants (Figure 5E, Figure S6) as well as in worms grown on these bacteria (Figure 5F, Figure S6). Despite functional rescue at the host level, and a similar growth profile to *dnaK* mutants alone (Figure S6A), the *dnaKclpX* double mutant exhibited a unique proteomic signature distinct from both single mutants and wild-type *E. coli* (Figure S6B), as well as in worms (Figure S6C). To capture the loss of *dnaK* bacterial effects by *clpX* on host UPS, we performed KEGG analysis in both *E. coli* (Figure S6D-H) and *C. elegans* (Figure S6J-M) with a focus on the comparison between up- and down-regulated proteins of *dnaKclpX* versus *dnaK* (Figure 5E-F). We observed a shared enrichment of central carbon and amino acid metabolism pathways, particularly branched-chain amino acids (BCAA) valine, leucine and isoleucine in bacteria (previously identified as an important UPS regulator^39^) and both BCAA and propionate metabolism in the host. BCAA degradation and propionate metabolism are two directly connected metabolic pathways through the sharing of the intermediate propionyl-CoA, further suggesting a role for microbe-host metabolic cross-talk in modulating host proteostasis. To determine how *E. coli* regulated BCAA and propionate metabolism in the host, we compared all proteins that were significantly down-regulated in *E. coli dnaK* mutants while up-regulated in both *clpX* and *dnaKclpX* mutants (Figure 5G). From the 33 proteins shown to be significant in these comparisons, the three proteins of the TonB-ExbB-ExbD energy transduction complex were strongly enriched for the Gene Ontology Molecular function of energy transducer activity (FDR=0.0111, Strength=2.1) which are involved in the transport of iron and vitamin B12 (VB12)^40^. Given the well described role of bacterial VB12 in the regulation of BCAA and propionate homeostasis in the host^32^, we investigated whether VB12 and/or propionate metabolites regulated UPS proteostasis. First, we created triple mutants of all known iron transporters in *E. coli* and observed that only the deletion of TonB significantly increased the levels of UPS fluorescence when compared to the effect observed when fed *dnaKclpX* mutants (Figure 5H, p=0.0008, Figure S6N) suggesting a regulation of TonB levels by DnaK that are controlled by ClpX. Consistent with this observation, the overexpression of TonB in a *dnaK* or *dnaKclpX* mutant significantly reduced UPS levels (Figure S6O). Overexpression of BtuB protein, a specific transporter of VB12, also reduced *dnaKclpX* levels to baseline levels (Figure S6P) and this required TonB, confirming that canonical tonB-BtuB-dependent VB12 transport is central to this regulatory axis.

**Figure 5.**
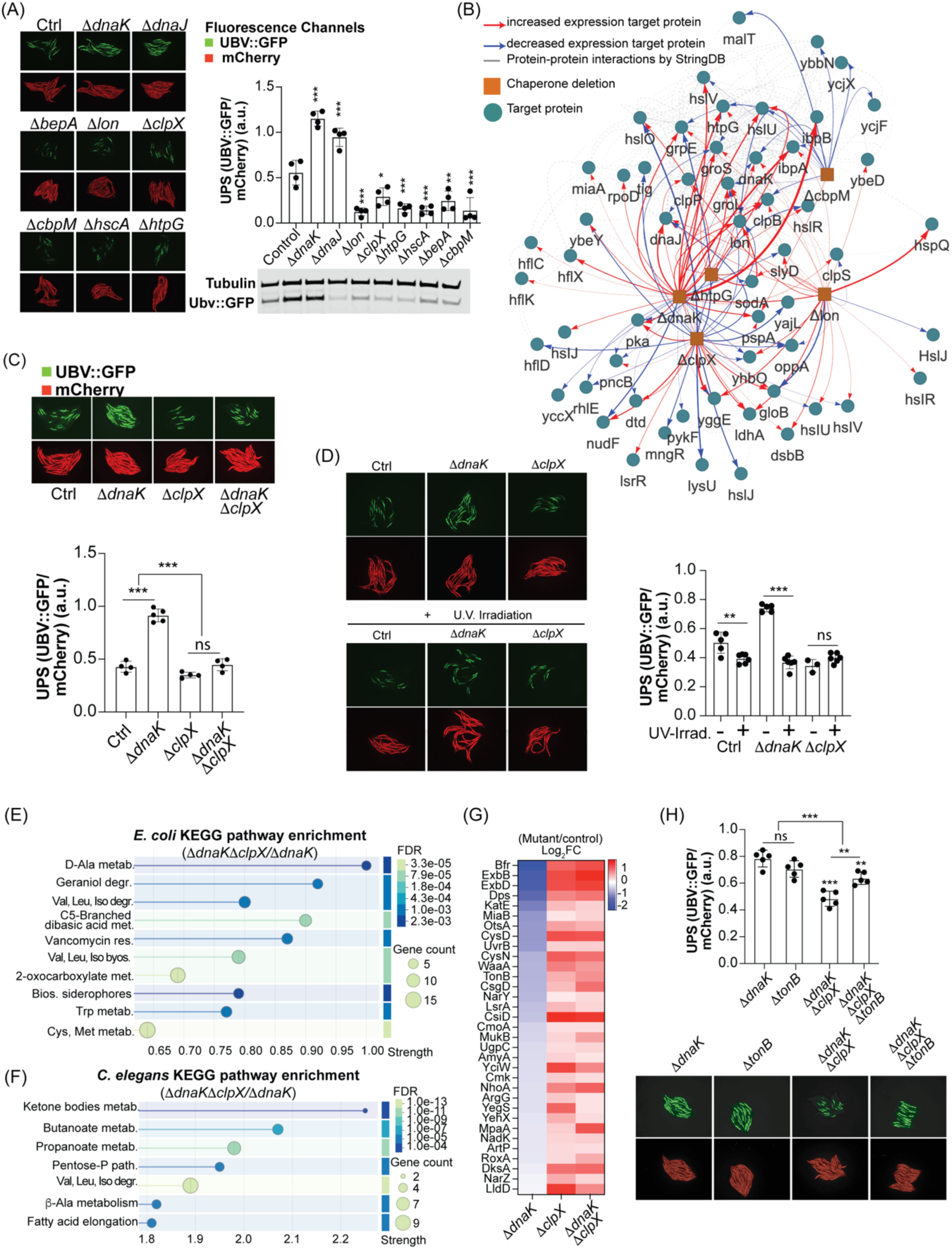
Proteostasis state in *E. coli* regulates proteostasis regulation at the host level. (A), Left, representative fluorescent microscope images of the worm reporters Ub(G76V)–GFP and constitutively expressed mCherry reporters fed on selected *E*. *coli* mutants identified in the Figure 4F bacterial proteostasis screen and respective quantification on the right. Western blotting of worm GFP and tubulin (housekeeping protein) for the selected proteins (n=4). (B), Bi-partite network representation of the proteome derived from proteomic analysis of the knockout *E. coli* genes Δ*dnaK*, Δ*lon*, Δ*clpX* and Δ*cbpM* compared to the control strain. Nodes are bacterial strains (orange) and significant proteins (grey). Edges represent protein expression (red for increased, blue for decreased) and protein-protein interactions from STRING (grey). (C), Representative images of normalized UPS fluorescence of worms fed on BW25113 and mutants Δ*dnaK*, Δ*clpX* and Δ*dnaK*Δ*clpX* and respective quantification (n=4). (D), Left, Representative fluorescence images from worms fed alive BW25113 (Control) and Δ*dnaK* and Δ*clpX* mutants cells (top) and UV-irradiated cells (bottom). Right, normalized UPS fluorescence quantification (n=5). (E,F) KEGG Pathways enrichment analysis from proteomics data comparing Δ*dnaK*Δ*clpX* double mutants to Δ*dnaK* single mutants obtained from *E. coli* cells only (E) or *C. elegans* grown on the respective mutants (F). (G), Heat map of the top significant proteins from proteomics data obtained from *E. coli* cells for the Δ*dnaK,* Δ*clpX* and Δ*dnaK*Δ*clpX* versus BW25113 comparisions. (H), Top, normalized UPS fluorescence quantification of worms fed *E. coli* mutants Δ*dnaK*, Δ*dnaK*Δ*tonB*, Δ*dnaK*Δ*clpX*, Δ*dnaKΔclpX*Δ*tonB* (n=5). Bottom, Respective representative fluorescence images of the Ub(G76V)–GFP and constitutively expressed mCherry reporters from the worm fed *E. coli* chaperone mutants Δ*dnaK,* Δ*tonB*, Δ*dnaK*Δ*clpX* and Δ*dnaK*Δ*clpX*Δ*tonB*. Stats correspond to comparison against the control (one-way ANOVA) and between nested conditions (two-way ANOVA), represented as *P < 0.05, **P < 0.01, ***P < 0.001, NS P > 0.05.

### Vitamin B12 dependent and independent flux regulates host UPS

To test the direct role of VB12, we supplemented VB12 and observed a uniform decrease of UPS levels in all bacterial mutant backgrounds (Figure 6A) but significant differences in the remaining UPS levels between them suggested the role of additional metabolites shaping host proteostasis. Given the role of VB12 in regulating propionate metabolism, we supplemented propionate and found that propionate increased UPS levels in worms fed control bacteria and *dnaKclpX* mutants but not *dnaK* (Figure 6B). Together with our proteomic data (Figure 5E), it suggested a role for propionate as a potential metabolite regulating host proteostasis. Deletion of the *sbm* operon, which encodes enzymes for the “sleeping beauty” mutase pathway that converts BCAAs (in particularly isoleucine) into propionate^41^, abolished the *dnaK*-induced UPS increase without affecting bacterial fitness (Figure 6C, Figure S6Q). These findings directly link bacterial propionate production to host UPS activation. Together, these data show for the first time that bacterial proteostasis modulates host proteostasis through metabolic signals.

**Figure 6.**
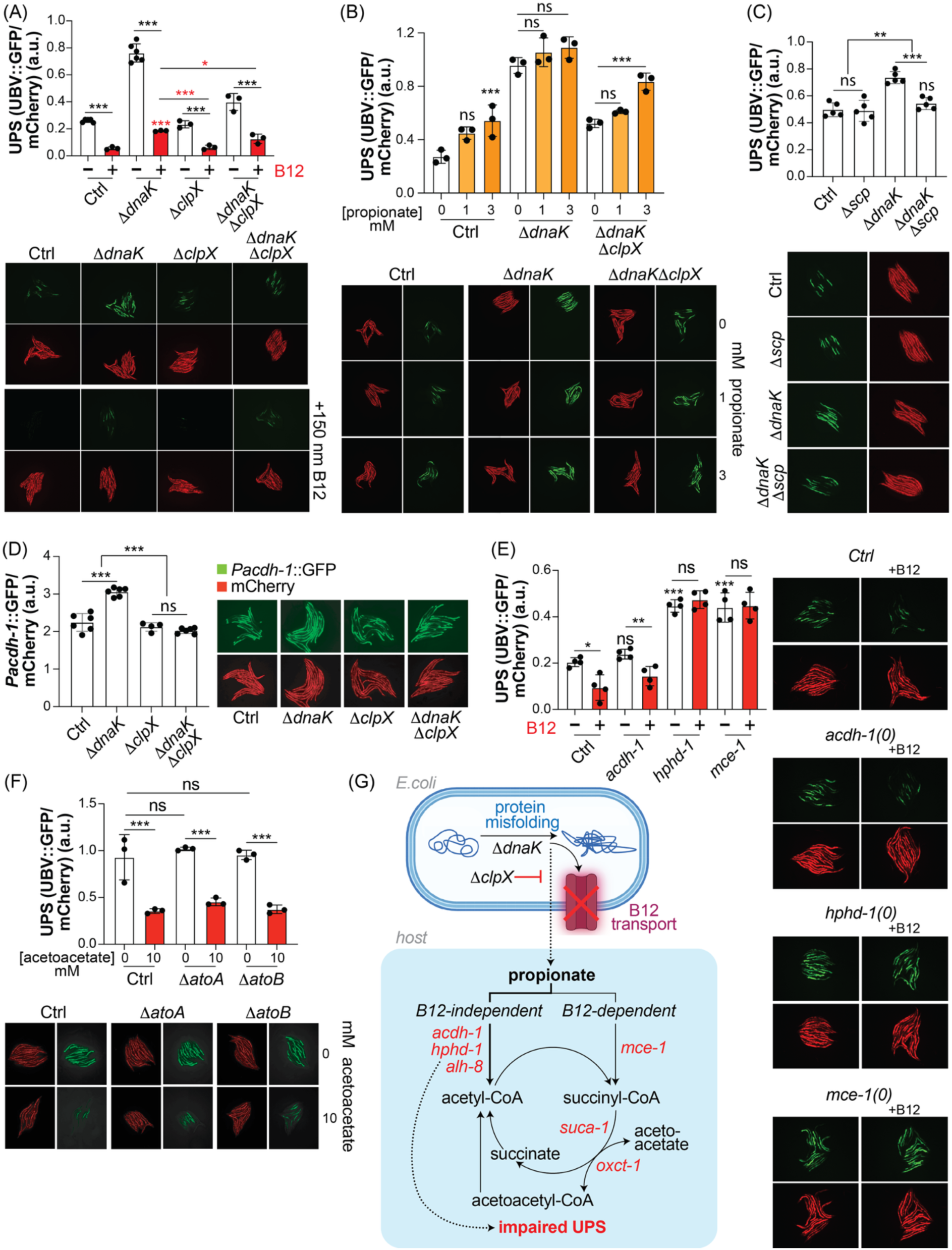
Vitamin B12 dependent and independent propionate catabolism regulates host proteostasis. (A), Top, normalized UPS fluorescence quantification of worms fed *E. coli* BW25113, Δ*dnaK*, Δ*clpX*, Δ*dnaK*Δ*clpX* in the presence (+) or absence (-) of vitamin B12 (150 nM) (n=3-6) and bottom, representative fluorescence images of the Ub(G76V)–GFP and constitutively expressed mCherry reporters. (B), Top, normalized UPS fluorescence quantification of worms fed on BW25113, Δ*dnaK* and Δ*dnaK*Δ*clpX* supplemented with propionate (0, 1 and 3 mM) (n=3). Bottom, representative fluorescence images of the Ub(G76V)–GFP and constitutively expressed mCherry reporters. (C), Top, normalized UPS fluorescence quantification of worms fed on BW25113 and strains Δ*scp*, Δ*dnaK*, Δ*dnaK*Δ*scp* (n=5). Right, representative fluorescence images of the Ub(G76V)–GFP and constitutively expressed mCherry reporters. (D), Letf, normalized UPS fluorescence quantification of worms fed on BW25113 and strains Δ*dnaK*, Δ*clpX* and Δ*dnaK*Δ*clpX* (n=4-6). Right, representative fluorescence images of the Ub(G76V)–GFP and constitutively expressed mCherry reporters. (E), Left, normalized UPS fluorescence quantification of worms fed on BW25113 for wild-type *C. elegans* N2 (Ctrl) and worm mutants *acdh-1(0)*, *hphd-1(0)*, *mce-1(0)* in the presence (+) and absence (-) of vitamin B12 (150 nM) (n=4). Right, representative fluorescence images of the Ub(G76V)–GFP and constitutively expressed mCherry reporters. (F), Top, normalized UPS fluorescence quantification of worms fed on BW25113 and strains Δ*atoA*, Δ*atoB* with and without acetoacetate (10 mM) (n=3). Bottom, representative fluorescence images of the Ub(G76V)– GFP and constitutively expressed mCherry reporters. (G), Diagram showing that proteostasis regulation at the bacterial level modulates the host response and proteostasis status via propionate and vitamin B12 dependent and independent pathways. Statistics correspond to comparison against the control (one-way ANOVA) and between nested conditions (two-way ANOVA), represented as *P < 0.05, **P < 0.01, ***P < 0.001, NS P > 0.05.

In *C. elegans*, propionate catabolism proceeds through a VB12-dependent canonical pathway and a VB12-independent “propionate shunt,” whose activation can be monitored by the *acdh-1p::GFP* reporter^32^. Worms fed *dnaK* mutants showed increased *acdh-1* levels, indicating elevated flux through the VB12-independent shunt, whereas worms fed *dnaKclpX* bacteria did not (Figure 6D). These findings are consistent with our proteomic evidence that *dnaKclpX* suppresses *dnaK*-driven metabolic rewiring. Genetic inhibition of the shunt downstream of *acdh-1* (e.g., *hphd-1, alh-8*) or the canonical pathway (e.g., *mce-1*) elevated UPS activity (Figure 6E, Figure S6R), while VB12 supplementation reduced UPS levels in wild-type and *acdh-1* mutants but not in *mce-1 or hphd-1* mutants. These findings demonstrate that the balance between VB12-dependent and -independent propionate catabolism determines the impact of propionate on proteostasis.

Proteomic analyses of worms fed *dnaK* versus *dnaKclpX* bacteria revealed enrichment of pathways linked to ketone metabolism (Figure 5F), aligning with previous reports that perturbations in propionate catabolism can alter ketone body pathways^42^. RNAi of *suca-1*, which contributes to conversion of acetoacetate to acetoacetyl-CoA (Figure S6R), increased UPS activity, whereas exogenous acetoacetate supplementation (independent of its degradation by *atoA* or *atoB*) decreased UPS levels (Figure 6F). This suggests that ketone intermediates can directly modulate host proteostasis. Collectively, these findings define a mechanistic chain (Figure 6G) in which bacterial DnaK–ClpX–TonB–BtuB control bacterial proteostasis and VB12 transport, thereby shaping propionate production and metabolic routing in the host. This balance between toxic VB12-independent catabolites and protective VB12-dependent flux, ultimately tunes UPS activity and proteostasis in *C. elegans*.

### DISCUSION

Metagenomic sequencing and other state-of-the art technical advances now enable high throughput, high-resolution scale analyses of microbial strains across diverse and complex ecosystems ranging from the human gut to marine and soil environments. Strain-level resolution has recently been shown to be crucial in microbiome research and in dictating microbe-host interactions. For example, strains of the same species can have diametrically opposed functional, ecological, and clinical manifestations, with species-level identification often leading to erroneous interpretations^43^. Strain-level characterization has also been emphasized in how bacterial strains are transmitted in human populations, highlighting the importance of the need to consider their biological effects ^44,45^. *Escherichia coli* has become a canonical example of the diversity displayed by a bacterial species, showing that its vast accessory genome harbored in its open pangenome contains an extensive array of bacterial functions that can potentially alter host physiology^15,19^. Our panel shows that any two given strains can differ by more than 50% of their genetic content. A central challenge in understanding the microbiome is reconciling the inherent genetic diversity contained within bacterial species and how this affects host physiology^46,47^.

Here, we present the most comprehensive analysis to date of how strain level variation within a single bacterial species shapes host responses. Given the challenge in defining bacterial functions for the poorly described microbial accessory genes, methods based in the transformers architecture have been trained over large protein databases creating physical representations of the protein space encoded in the microbiome^48,49^. These latent spaces have proven to be an accurate representation of protein function and have been used recently to capture the vastly and highly uncharacterized defense systems in bacteria ^22,50^. In this work, we leveraged the protein embeddings predicted by the pLM^21^ to study how the latent functional landscape encoded in the *E. coli* pangenome regulates host physiology. By compressing the genomic and functional information encoded in thousands of *E. coli* strains into unified “strain embeddings”, we created a geometric map that captures the potential function per strain, which can be mapped onto the host phenotype it elicits. By building a high-throughput panel of strain-host mono-association spanning hundreds of *E. coli* strains, we have established an experimental platform validating our computational approach and providing additional mechanistic insights. Together, our data uncovers several fundamental principles in microbe-host biology at the strain level. 1) Each strain elicits a unique molecular signature in the host; 2) our data shows that the protein functions shared by every strain (the core) are located in a narrow geometrical space compared to the vast strain-specific functions, 3) strains within the same phylogroup display an ∼60:40 of positively to negatively correlated effects on host responses; and 4) simple phylogenetic relatedness between strains does not predict the host molecular programs they induce. Our data supports the hypothesis that phylogeny, while important, is inadequate as a single factor to link bacterial functions to host physiology. Future work will be required to determine how strain-level effects manifest in the context of complex microbiota and to test whether these basic principles extend to other bacterial species with open or closed pangenomes.

In line with this, the canonical laboratory microbial source for *C. elegans*, the *Escherichia coli* OP50 strain, elicits a distinct and divergent molecular response in the host compared with other *E. coli* strains (Figure 1E). This observation is consistent with a growing number of studies incorporating multiple bacterial strains reporting strain-dependent mechanisms underlying diverse host phenotypes, including drug responses ^51,52^, behavior, reproduction, and lifespan ^31,53,54^. Collectively, these findings suggest that the experimental convenience afforded by *E. coli* OP50 may be offset by the specific molecular and physiological signature it imposes on *C. elegans*, potentially failing to reflect the “true” wild-type biology of the host and motivating a critical re-evaluation of the foundational literature of an entire field. This may possibly be better captured using native microbiome *C. elegans* strains or alternatively, commensal *E. coli* strains employed in this study. Using a protein-embedding framework, this work supports these claims as it identifies a broad repertoire of bacterial functions, spanning many COG functional categories with known effects on host physiology, that are regulated at the level of the bacterial pangenome. Among the most significantly enriched categories is proteolysis. Notably, recent work has demonstrated that differences in bacterial-derived RNAs between *E. coli* OP50 and HT115 can trigger a systemic response in *C. elegans* that protect against protein aggregation during aging^55^. Likewise, the present study reveals pronounced differences among *E. coli* strains (Figure 3E) and strains belonging to bacterial species from other phyla (Figure S4F) in their ability to modulate the host ubiquitin–proteasome system. We demonstrate that key bacterial proteostasis regulators control host UPS activity by modulating the availability of vitamin B12 and propionate, which in turn dictates the flux through host propionate degradation pathways. Bacteria that produce or efficiently scavenge B12 can control community composition and metabolic activity by outcompeting B12-dependent neighbors ^56,57^. For example, B12 production by *Eubacterium hallii* enables *Akkermansia muciniphila* to convert succinate to propionate, shifting succinate levels, and thereby reshaping the surrounding metabolic network ^58^. Here, an unanticipated mechanism is described in which the DnaK/J chaperone system and the ClpX protease act in concert to fine tune B12 and propionate levels. While this regulatory axis may have evolved primarily to modulate microbial community interactions, it unexpectedly alters host proteostasis, giving rise to what can be conceptualized as microbe–host “co-proteostasis” derived from microbe-host co-metabolism cues.

## METHODS

### RNA sequencing of *C. elegans* fed on PG *E. coli* strains

Around 30 *C. elegans* animals were grown per well in 96-well microtiter plates, each well containing NGM seeded with a distinct *E. coli* strain. On day 1 of adulthood the worms were transferred in a high-throughput manner to clean 96 well plates using INTEGRA Viaflo 96 liquid handler. Worms were washed twice with RNase-free water to remove bacterial traces and flash-frozen in liquid nitrogen. For lysis, we used bead-based mechanical disruption (Bertin Technologies) in RNA lysis buffer (Zymo Research) and on a bench top Eppendorf Thermomixer C at 2000rpm for 20 min at 4°C. RNA was concentrated and purified with the RNA Clean & Concentrator-96 kit (Zymo Research). Samples were eluted into microtiter plates and stored at -80°C prior to library preparation. We quantified and checked the integrity of the RNA and selected batches of 48 RNA samples with similar quality to ensure uniform RNA input. To obtain comprehensive coverage of expressed genes in the *C. elegans* host, we employed Lexogen QuantSeq-Pool Sample-Barcoded 3’ mRNA-Seq Library Preparation. Each RNA sample was labelled with a unique 12-nucleotide i1 sequence barcode before conversion to cDNA and pooling. Before amplification, each cDNA pool was dual-indexed with 12-nucleotide i5/i7 index sequences. To validate the RNA extraction and library preparation we prepared a test-pool library from conventional *E. coli* laboratory strains. We performed pair-ended sequencing of the test pool on an Illumina MiSeq sequencer and obtained successful demultiplexing. In total, we prepared 16 libraries that were normalized for final pool sequencing based on library quantification by Qubit 3.0 fluorometer and average library size measured by TapeStation 4200. To remove *E. coli* phylogroup representation biases we randomly distributed strains in the different libraries.

Sequencing was performed on an Illumina MiSeq. Sequences were demultiplexed using DRAGEN GenerateFastQ (v3.7.4) by using the i5/i7 barcodes to separate the different libraries. Each library was further demultiplexed by using idemux (v.0.1.6) and by using each library and sample i1 barcode identifiers. Sequences were quality-cleaned with trimmomatic (v.0.39), removing Illumina adapters and dropping sequences below 65 nucleotides. Lexogen recommends trimming the first 12 nucleotides of each read, a step that can be avoided if the aligner used to map the reads can perform soft-clipping, which was our case with salmon (v.1.10.1). Outlier samples with extremely low read count were discarded at this point. Quality-filtered reads were then filtered to exclude samples with less than 4x10^5 reads, resulting in the discard of 59 samples mainly from libraries 14 and 15. 661 samples were kept for posterior analyses (606 unique strains). Reads have been mapped to the cDNA of the *C. elegans* reference genome in Ensembl (version 111) with salmon (v.1.10.2), which performs soft-clipping, to build the counts matrix and then imported to R with tximport (v.1.28). were analyzed with DESeq2^59^ (v.1.40.1) using the phylogroup as the main group and using the library information to avoid possible batch effect Genes that had less than 10 reads in total were discarded. PCA calculations were performed with the plotPCA function from DESeq2 with the data transformed with the vst function. Outliers from the PCA plot from normalized counts but not batch-corrected were removed from the analysis. Batch correction was performed with the removeBatchEffect function from limma^60^ (v.3.64.1). Per-gene variance was modelled with the function modelGeneVar from the scran package^61^ (v.1.36.0).

To build the transcriptional landscapes from the worm, the curated database from WormCat^23^ containing categories at level 1 to 3 was downloaded (Nov-2021 version). The genes belonging to each category per level were summed by using the normalized and batch-corrected reads from the transcriptional profile. A single value was obtained per category, level and strain, which was used for downstream analyses.

### *E. coli* strain landscape selection

The *E. coli* strains were selected from the NCBI genome database. The metadata was downloaded on January 11th, 2024, and was downloaded with the NCBI dataset download tool (v. 16.2.0). The strains were filtered based on the criteria described here. Genomes that were not at the assembly level of “chromosome”, “complete genome” or “scaffold” were removed. The scaffold N50 was used to filter out genomes with a value lower than 150K. CheckM metrics were used to remove genomes with a completeness lower than 95% and a contamination higher than 1%. Mash distances (mash v. 1.1)^62^ were calculated pairwise between all genomes, and those strains whose mean distances were higher than 0.05 were removed. Finally, genomes with a sequence length greater than 7Mbp, and/or genomes with a contig count higher than 300 were removed. This resulted in 8,829 assemblies passing all filters. 517 strains were added from the EcoRef collection^19^, where evolution-related strains were discarded from the analysis. 212 commensal strains were added from human isolates from Australia^15^. Phylogroups were assigned with the EzClermont v. 0.7.0 tool^63^ (tool based on the approach from ClermonTyping^64^), and genomes belonging to class cryptic, U/cryptic and fails were discarded. The final number of genomes was 9,558 assemblies.

### Gene annotation and pangenome analysis

Genome annotation was performed with Bakta^65^ (v. 1.9.3) using the full database (v. 5.1) using by default parameters. The pangenome was analyzed with Panaroo^66^, selecting a strict clean mode and removing invalid genes. Due to the complexity of this pangenome and the computation time, the pangenome was split into 5 parts containing approximately 2000 random genomes each. Each pangenome was calculated using the same parameters. The output from the 5 calculations was merged into the final output using the Panaroo-merge function from the main pipeline. The reference sequences from the gene families were then translated into proteins (using a custom Python script with biopython v. 1.84). Gene presence/absence matrix was used to calculate the Principal Component Analysis shown in the main text by using the PCA function from scikit-learn v. 1.5.1. The pangenome analysis from the strains used for the *C. elegans* transcriptome and the reporter screening was performed independently by leveraging the annotations obtained with Bakta and running the pipeline as a single process this time (592 strains in total).

Gene presence/absence matrix was used to generate the accumulation curve (ACC) for the full *E. coli* panel of 9,558 strains, and to calculate the Heap’s law. The ACC was generated by first removing gene families that were present in more than 99% of the strains, then dividing the total gene count into 50 sampling points and randomly picking genomes for each sampling point to count the number of genes. This process was iterated over 5 times. Heap’s law was calculated to fit the following equation:

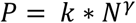

P is the pangenome size, N is the number of genomes, and *k* and *γ* are the parameters to fit. Heap’s law parameters were fitted using the average of the ACC data per point with the nls function in R.

Pairwise strain genetic similarity was measured as the Jaccard similarity between every pair of strains included in the study. It was calculated as the shared genetic content divided by the union of their genetic content between each strain pair.

### Phylogeny

From the core genome extracted from Panaroo, a subset of 275 genes present in all strains was used to build the phylogenetic tree for the full *E. coli* panel. To avoid multicopy bias, only one gene per strain was kept for the alignment. For the laboratory strains included in the smaller panel we used the full core genome. The alignment was done using mafft^67^ (v. 7. 526), and the tree was constructed with IQ-Tree^68^ (v. 2.3.6) with the GTR+I+G substitution model.

### GO term prediction

#### Sequence-based methods

Proteins were classified using two of the most popular sequence-based methods used in the community: InterPro and eggNOG. Search in the InterPro databases was done using interproscan v. 5.59-91.0^26^ with by default parameters. The search in the eggNOG database was done using eggnog-mapper v. 2.1.12^27^ with MMseqs2 to look for novel families options enabled. Results from both methods were filtered to remove entries that had an E-value larger than 1e-5.

#### Machine learning methods

Reference genes from Panaroo were split into 4 smaller files to fit in memory. Proteinfer^28^ source code was downloaded from github and function prediction was done by using 5 ensemble models and a reporting threshold of 0.3.

The reference genes from Panaroo were translated into proteins and sequences were clustered with CD-HIT v. 4.8.1 (similarity threshold of 0.98) to remove similar sequences from the dataset, resulting in 55,942 unique clusters. The resulting file was split into 20 smaller files to fit into memory. Proteins were embedded with bio-embeddings pipeline (v.0.2.2), by using the model ProtT5-XL-U50^21^ in half-precision mode. Proteins larger than 3,000 amino acids were discarded to fit in memory. Transfer learning was done using available pipelines under bio-embeddings that used goPredSim^29^, using Euclidean distances and a k-nearest-neighbors of 3. ProtT5 h5 file was used as a reference with GOA annotations from 2022. Proteinfer, protein embeddings and transfer learning were carried in a computer with 32Gb of RAM and an RTX 4080 GPU with 16Gb of memory.

#### Information content calculation

To calculate the information content (IC) of the GO terms predicted by the different tools, we used an adaptation of the method from Barrios-Nuñez *et al*^25^. Given that GO terms have a hierarchical structure, the deeper nodes from the branch will contain a higher functional information. Considering that having a deep node in the branch is less likely than to have a higher node with less information, we can approximate the information content of each node by the negative logarithm of the probability for that node to be inferred:

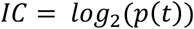

Where p(t) is the probability for that node, which can be calculated as:

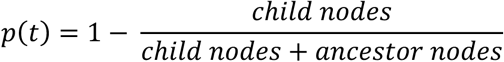

IC was calculated by joining all the GO term predictions together to create a joint library of terms for the pangenome. Given that we lack a pre-computed list of GO terms with their probabilities as exist for reference organisms, we had to calculate these probabilities from scratch. We joined together all GO term predictions from the 4 methods and kept the uniquely present GO terms. This allowed us to create a database whereby to filter the resulting steps. To calculate the number of ancestors and child nodes from each term, OWLTools was used (release 2024-06-12). The database used is the go-basic.obo from geneontology.org (accessed in October 2024). From the joint set of unique GO terms we used the OWLTools-Runner function to get both the ancestors and descendants from each node. As the descendants from a node, especially from the ones up in the tree, can have many different child nodes depending on the final function, we removed all the GO terms that were not present in our joint dataset. The probability was calculated as defined but corrected as 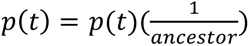 for the cases where no descendant was kept in the list, but the GO term did not reach the bottom of the branch from the obo database.

#### Strain embeddings calculation

Strain embeddings were calculated based on the gene presence/absence matrix generated by Panaroo, the protein embeddings generated by the pLM model ProtT5-XL-U50, and the number of genes per strain. To calculate any of the different strain embeddings versions described below, we excluded the core genome set of genes, as they were not useful given that all strains shared them.

Strain embeddings were generated using three distinct aggregation methods: 1) direct summation via matrix multiplication; 2) simple averaging, normalized by the gene count per strain to mitigate genome size bias; 3) weighted averaging, which employs an Inverse Gene Frequency (IGF) metric. The three versions can be visualized in Figure S S3a, the strain embeddings have been uploaded to Zenodo (https://doi.org/10.5281/zenodo.18221759)

#### Matrix multiplication

The simplest form was calculated by multiplying the presence/absence matrix with the embedding matrix with the following form:

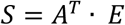

Where A is the binary matrix of gene presence/absence with *nxm* dimensions (genes and genomes), E is the matrix of protein embeddings from the representative genes with *nxd* dimensions (genes and embeddings), and S is the objective strain embeddings with *mxd* dimensions (genomes and embeddings).

#### Average strain embeddings

The average strain embeddings were calculated based on how many genes were encoded in each genome and then applying a diagonal normalization on the matrix multiplication equation. The diagonal normalization is a *mxm* matrix where the diagonal is the inverse of the number of genes per strain, where *N_i_* is the number of present genes in strain *i*:

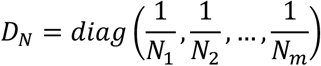

Therefore, the average strain embeddings were calculated as:

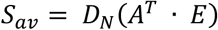

Where *S_av_* is the objective average strain embeddings with dimensions *mxd*.

#### Weighted average strain embeddings

Finally, the contribution for each gene to the strain potential was scaled in terms of their proportion, thus, increasing the importance of rare genes to the final position of the strain embedding. That is, genes that are common have a lower weight than the ones that are rarer. To do so, relied on an adaptation of the *Inverse Document Frequency* metric that can be adapted here as the *Inverse Gene Frequency* (IGF).

First, the Strain Count for a gene family (*C_i_*) was defined as the number of strains in which the gene family *i* was present over the total number of strains (*M*). This was equivalent to the sum of the *i*-th row of matrix A:

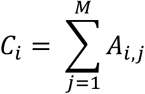

We next defined Weight for Gene Family *i* (*W_i_*) as the logarithm of the relative presence of a specific gene family, where *W_i_* = 0 if *C_i_* = *M* (gene present in all strains):

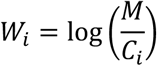

We then defined the diagonal matrix with gene weights calculated from last equation as:

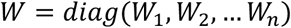

Here *W* is a matrix of *nxn* dimensions. We then used this matrix to calculate the weighted protein embeddings (*E*_W_) as:

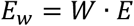

Where *E*_W_ and *E* are matrices with *nxd* dimensions. Then we calculated the weighted sum of embeddings (*S*_Weighted_*sum*_) for each strain as:

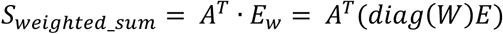

Where *S*_Weighted_*sum*_ is a *mxd* matrix. We proceeded by calculating the sum of weights for each strain (*W_sum_*) as:

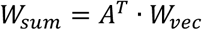

Where *W_vec_* is an *n x* 1 column vector containing the *W_i_* values; and the sum of weights for each strain *j* is 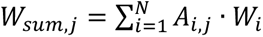

We created another diagonal normalization matrix (*D*_W_, an *mxm* matrix), where the diagonal elements are the inverse of the sum of weights for each strain:

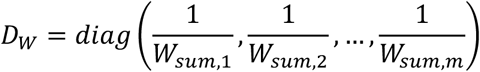

Finally, we used all these outputs to do the final calculation and got the weighted averaged strain embeddings:

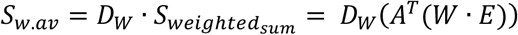

Where *D*_W_is a *mxm* matrix, *S*_Weighted_*sum*_ is a *mxd* matrix, and the product *S*_W.*av*_ is a *mxd* matrix whose each vector row *S*_j_ is the weighted averaged embedding for strain *j*.

#### Max-pooling strain embeddings

To capture the most prominent functional features present within a strain’s accessory genome regardless of gene background abundance, max-pooled strain embeddings were generated. Given the binary gene presence/absence matrix *A* (where *A_i_*_,j_ = 1 if gene family *i* is present in strain *j*) and the protein embedding matrix *E* containing ProtT5 representation vectors 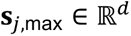 for strain *j* is calculated as:

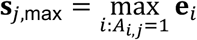

where the maximum operation is performed element-wise across all *d* embedding dimensions for all gene families present in strain *j* (*A_i_*_,j_ = 1).

#### Generalized mean strain embedding

To establish a flexible pooling framework capable of interpolating between average pooling and max-pooling, Generalized Mean Pooling (GeM) was implemented. GeM introduces a parameter *p* that controls the degree to which prominent functional features dominate the final strain vector. For strain *j* with 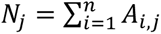 present gene families, the GeM strain embedding vector 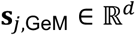 is defined as:

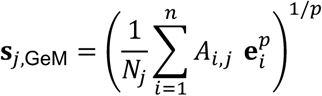

At *p* = 1, GeM is equal to standard average pooling, whereas higher values of *p* progressively approaches to max-pooling as *p* → ∞. GeM pooling representations were evaluated across *p* = 2, 4, 10 to test intermediate steps between the standard average pooling and max-pooling strain embeddings.

#### Functional mapping onto host phenotype

Average strain embeddings from the 9,558 *E. coli* strains were used to create a Principal Component Analysis in R using the prcomp function. The PCA coordinates were then leveraged to create the pangenome-host functional mapping. The WormCat-aggregated functions at level 3 were mapped onto the PCA coordinates 1 and 2 of the laboratory *E. coli* strains. The worm functions were correlated to each Principal Component per separate by using Spearman correlation and by correcting the P-value for multiple comparisons with a FDR calculated with the Benjamini-Hochberg method.

#### High-throughput imaging and data analysis

*C. elegans* animals were synchronized by standard hypochlorite method, and around 20 L1 worms were transferred to each well of 96-well plates seeded with *E. coli* pangenome strains. The worms were incubated at 20°C until D1 of adulthood and immobilized for imaging with 5 µL of 2% levamisole per well using INTEGRA Viaflo-96. Images were acquired by an automated protocol that captured 10 images at fixed z-heights per well under identical exposure settings using a Zeiss Axio Zoom V16 microscope system equipped with an AxioCam camera operated by Zen 2 software (Zeiss). GFP filter set (excitation: 450-490 nm; emission: 500-550 nm) or the RFP filter set (excitation: 559-585 nm; emission: 600-690 nm) was used depending on the strain being imaged. All images were exported in CZI format, and the most focused z-stack was extracted in Python (v. 3.12). Ilastik (v. 1.4) was used to detect worm pixels and to quantify fluorescence levels per worm/cluster of worms.

The fluorescence data was then filtered such that only single worms (ilastik single worm probability > 0.5) with a pixel size between 1000 and 6000 or clustered worms (ilastik clustered worm probability > 0.5) with a pixel size greater than 6000 were retained. Worm mean fluorescence expressed as the brightness per worm as a whole and corrected per size, was corrected against the background for each case. The mean-fluorescence of technical replicates was then averaged, and biological replicates were normalized. Mean-fluorescence was normalized such that, for each worm reporter, the median of each biological replicate was equal to the global median of all biological replicates. Following this, for each worm-bacteria pair the biological replicates, where n > 2 (up to n=5), were subjected to a z-score analysis using scipy.stats.zscore module (SciPy v1.8) and biological replicates with an absolute z-score > 1.5 for mean-fluorescence were removed. Given that most data had only 2 biological replicates and further replicates were only performed in select cases to replace replicates where some wells reduced data quality, from these sets of biological replicates the two replicates with the lowest deviation from each other were carried forward. Where only 1 biological replicate (8.9% of worm-bacteria pairs) was available; these were carried forward alone. Single replicates arise due to empty wells, where reporter worms escape wells and do not appear in the images; however, differences between single replicates and double replicates were broadly inexistent. The log2 ratio between each biological replicate pairs’ mean-fluorescence value was then calculated, and the median log2 ratio calculated for each reporter worm. For each reporter worm dataset, a decreasing threshold was iteratively tested for the maximum allowed log2 ratio between replicates and the maximum deviation from the median log2 ratio. Here, data above the threshold was removed and the Pearson correlation between the biological replicates calculated using scipy.stats.pearsonr module (SciPy v1.8). The threshold was continuously decreased and worm-bacteria pairs removed until a Pearson correlation of >= 0.7 was achieved. The final mean-fluorescence values were then calculated from the average of the biological replicates. Values were then normalized against the median for each reporter case for the tree representation, which was visualized with tidytree (v. 0.4.6), treeio (v. 1.32.0) and phytools (v. 2.4-4).

Pearson correlations of median ratio profiles were calculated for all strain-strain pairs to produce a correlation matrix using pandas.DataFrame.corr (pandas v2.1.4). A phylogenetic distance matrix for strains was hierarchical clustered, using the UPGMA algorithm, to produce a linkage matrix. Hierarchical clustering was performed here using the scipy.cluster.hierarchy.linkage module (SciPy v1.13.1) and prior to clustering the distance matrix was converted from the vector-from to the square-form using scipy.distance.squareform (SciPy v1.13.1). The median ratio correlation matrix was then clustered either using the phylogenetic linkage matrix or by strain-strain correlation profile similarity and displayed as a clustered heatmap (Seaborn v0.13.2, matplotlib v3.10.3).

Pearson correlations of median ratio profiles were then calculated for all strain-strain pairs within the same phylogroup, as above. The percentage of strains with positive or negative correlations within each phylogroup, as well as for the pangenome, were calculated for a range of thresholds between 0 to 1 using steps of 0.05, excluding same-strain pairs. Where correlations were greater than the threshold, they were classed as positively correlated. Where correlations were less than the negative of the threshold, they were classed as negatively correlated. Clustered heatmaps were produced for each phylogroup correlation matrix, hierarchically clustering by strain-strain correlation profile similarity (Seaborn v0.13.2, matplotlib v3.10.3). Chord plots were calculated as the pairwise Pearson correlation and P-values corrected by Benjamini-Hochberg. The significance threshold was set at an alpha of 0.05, and results were represented as the symmetrical relation of the significant correlations existent per phylogroup with the library circlize (v. 0.4.16)

#### Microbial Genome Wide Association Study (mGWAS)

Microbial genome-wide association testing was performed to identify bacterial sequence variants associated with C. elegans transcriptional and reporter phenotypic responses. Sequence unitigs were generated and extracted from the *E. coli* pangenome assemblies using the microGWAS pipeline^69^ (v.0.5.1). Association testing between unitig presence/absence patterns and continuous host phenotypic scores across the strain panel was executed using pyseer (v. 1.4.0) using max-dimensions to 5. To control for bacterial population structure, linear mixed models were fitted using a genomic distance matrix derived from the core phylogeny. Association results were filtered using a dual-threshold strategy: a hard significance cut-off of − log_10_(*p*) ≥ 4.5 and a soft filter based on repeated-value frequency thresholds to eliminate structural biases and false positives caused by overrepresented unitigs. Significant unitigs passing both filters were mapped back to reference gene families for functional annotation using genomes reannotated from the panaroo results.

#### Permutational Jaccard Similarity Testing

To evaluate genetic overlap among bacterial drivers influencing distinct host biological processes, pairwise Jaccard similarity coefficients were calculated between the sets of significant mGWAS gene markers identified across host functional categories:

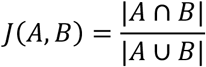

Where *A* and *B* represent the sets of mGWAS-significant *E. coli* gene families associated with two distinct host functional categories. To determine whether observed genetic overlaps were statistically higher than expected by chance, a permutation test was executed with 10,000 iterations. For each pairwise comparison, genes were pooled into a combined matrix and randomly redistributed into pseudo-categories matching the original set sizes to construct an empirical null distribution of Jaccard scores. Empirical *p*-values were computed as the proportion of permuted similarity scores meeting or exceeding the observed index and corrected for multiple testing using the Benjamini–Hochberg (BH) false discovery rate method (*p*_adj_ < 0.05).

Significant category pairs were assigned to broad functional domains based on the WormCat level 1 categories (Proteostasis, Stress, Metabolism, Cytoskeleton, and Cell Cycle). To map functional interconnections between host pathways driven by shared bacterial loci, both domain_1_ to domain_2_, and domain_2_ to domain_1_ were considered and represented with ggalluvial (v.0.12.6) in R. Category-level co-occurrence graphs were represented using tidygraph and ggraph (v.2.2.2) via a Kamada–Kawai force-directed layout.

#### Protein-Protein Interaction Network Reconstruction

Bacterial genes implicated in the proteostasis-1CC metabolism from the permutational Jaccard analysis were extracted for functional network mapping. Protein-protein interaction (PPI) data were retrieved from the STRING database using *E. coli* K-12 (MG1655) as the reference background. Network layouts were generated using ggraph (Kamada–Kawai layout), with edge width and color scaled to the STRING combined confidence score. To highlight key bacterial effectors within the network, node size and color fill were mapped to the mGWAS squared effect size (*β*^2^) derived from linear mixed model associations. Gene labels were selectively displayed for nodes exhibiting strong standardized effect sizes (*β* ≥ 0.25).

#### Machine Learning Predictive Benchmarking of Pooling Methods

To systematically rank and evaluate the ability of the seven strain embedding pooling representations (Addition, Average, Weighted Average/IDF, Max-pooling, and Generalized Mean Pooling (p = [2,4,10]) to preserve pangenomic functional features that drive host physiology, a high-throughput machine learning workflow was executed within the tidymodels framework (v.1.5.0) in R. Models were trained across 460 *C. elegans* functional category targets derived from WormCat Level 3 classifications.

Predictor matrices representing each pooling strategy were linked with target host phenotypes and bacterial strain metadata. Predictor columns were centered and scaled (step_normalize), and invariant variables were removed (step_zv). Model performance was evaluated using a 3-repeated 5-fold cross-validation scheme. Resampling splits were explicitly stratified by bacterial phylogroup.

Random Forest regression models were specified using parsnip (v.1.5.0) via the ranger computational engine. Each forest was constructed with 500 trees, and permutation-based feature importance was recorded. Hyperparameter space search for the minimal node size parameter was executed across 10 grid candidate states per workflow using workflowsets. Out-of-sample performance across hyperparameter configurations was evaluated using the coefficient of determination (R^2^). The top tuning state per pooling-workflow combination was retained to rank the predictability of host biological subsystems.

#### Proteostasis strains and culture conditions

Bacterial strains generated in this study and primers used are listed in the Table S12. *E. coli* BW25113 single gene deletion mutants were obtained from the KEIO collection and confirmed by PCR. The reaction was performed with GoTaq mix and the PCR was carried out in a PCRmax Alpha Cycler 2 as follows: 2min at 98°C for the initial activation of enzymes, 30 cycles of 30s at 98°C, 30s at 58°C and 1 min/Kb at 72°C. Each strain was grown in LB broth overnight and 120 μL were plated on nematode growth medium (NGM) plates and kept at 20°C for 2 days.

The *C. elegans* UBV reporter, PP607 (hhIs64[unc-119(+); sur-5::UbV-GFP]; hhIs73[unc-119(+); sur-5::mCherry]) was provided by Hoppe Lab, Germany. This strain allows to quantify the proteasomal activity *in vivo* thanks to the GFP fused to a non-cleavable ubiquitin (UbV-GFP) under the control of the ubiquitous *sur-5* promoter^30,35,36^. The following strains were made for fluorescence studies: FGC72 *nIs470[Pcysl-2::GFP];wbmIs67 [eft-3p::3XFLAG::wrmScarlet::unc-54 3’UTR *wbmIs65];* FGC73 *agIs17 [myo-2p::mCherry + irg-1p::GFP] IV;wbmIs67 [eft 3p::3XFLAG::wrmScarlet::unc-54 3’UTR *wbmIs65]*; FGC74 [rtIs30(pfat-7::GFP);wbmIs67 [eft-3p::3XFLAG::wrmScarlet::unc-54 3’UTR *wbmIs65]; FGC76 *wwIs25[Pacdh-2::GFP + unc-119(+)*];*wbmIs67 [eft-3p::3XFLAG::wrmScarlet::unc-54 3’UTR *wbmIs65]*; FGC77 *dvIs19 [(pAF15)gst-4p::GFP::NLS] III;wbmIs67 [eft-3p::3XFLAG::wrmScarlet::unc-54 3’UTR *wbmIs65];* FGC78 *agIs219 [T24B8.5p::GFP::unc-54 3’ UTR + ttx-3p::GFP::unc-54 3’ UTR] III;wbmIs67 [eft-3p::3XFLAG::wrmScarlet::unc-54 3’UTR *wbmIs65]*; FGC79 *wwIs24 [Pacdh-1::GFP + unc-119(+)];wbmIs67 [eft-3p::3XFLAG::wrmScarlet::unc-54 3’UTR *wbmIs65]*; FGC80 *zcIs13[hsp-6::GFP];wbmIs67 [eft-3p::3XFLAG::wrmScarlet::unc-54 3’UTR *wbmIs65]*; FGC81 *dvIs70 [hsp-16.2p::GFP + rol-6(su1006);wbmIs67 [eft-3p::3XFLAG::wrmScarlet::unc-54 3’UTR *wbmIs65];* FGC82 *mgIs73 [cyp-14A4p::gfp::cyp-14A4 3’UTR + myo-2p::mCherry] V.;wbmIs67 [eft-3p::3XFLAG::wrmScarlet::unc-54 3’UTR *wbmIs65];* FGC83 *muIs84 [(pAD76) sod-3p::GFP + rol-6];wbmIs67 [eft-3p::3XFLAG::wrmScarlet::unc-54 3’UTR *wbmIs65];* FGC84 *wuIs177 [Pftn-1::gfp lin-15(+)];wbmIs67 [eft-3p::3XFLAG::wrmScarlet::unc-54 3’UTR *wbmIs65]*; FGC89 *acdh-1(ok1489), hhIs72[unc-119(+); sur-5::mCherry], hhIs64 [unc-119(+); sur-5::UbiV-GFP]III*; FGC120 *hphd-1(ok3580); hhIs72[unc-119(+); sur-5::mCherry], hhIs64 [unc-119(+); sur-5::UbiV-GFP]III*; FGC121 *mce-1(ok243) I; hhIs72[unc-119(+); sur-5::mCherry], hhIs64 [unc-119(+); sur-5::UbiV-GFP]III*. Detailed information can be found in Table S11. Worms were maintained at 20°C, on nematode growth medium NGM seeded with different bacterial strains. We supplemented NGM with homocysteine (final concentration 1 and 5 mmol/L), propionate (final concentration 1 and 3 mmol/L) and cobalamin (vitamin B12, final concentration 150 nmol/L) solubilized in water and filter sterilized.

#### Double deletion bacterial strain construction

Double gene deletion has been generated by using strains from the KEIO Library^70^. This library is based on the *Escherichia coli* strain BW25113. The Kanamycin cassette has been removed by using the plasmid pCP20. This plasmid encodes the yeast Flp recombinase gene, chloramphenicol and ampicillin resistant genes, and temperature sensitive replication^71^. *E. coli* BW25113 strains containing a single mutation were transformed following the TSS enhanced chemical transformation^72^ and were plated on chloramphenicol 30ug/ml incubated at 30°C overnight. Clones were selected and streaked on LB with no selection and LB-Kanamycin (50 μg/mL) plates, incubated at 30°C overnight. Kanamycin sensitive clones were streaked on LB agar plates and incubated at 37°C overnight to stop the replication of the pCP20. Clones were then streaked on LB, LB-Kanamycin and LB-chloramphenicol and incubated at 37°C overnight. Sensitive clones to chloramphenicol and kanamycin were selected and kanamycin cassette removal was confirmed by PCR. From this, we obtained mutant with a single mutation not carrying a kanamycin cassette. The secondary mutations were then extracted from another mutant of the Keio library. The strain of interest was lysed by using P1 phage and transduced in *E. coli* kanamycin sensitive strain according to the protocol from Thomason *et al*. 2007^73,74^ and was then selected for his resistance to kanamycin. Finally, the presence of both mutations was confirmed by PCR.

#### Bacterial growth assay

The optical density (OD) at 595nm was monitored using NuncTM 96-well polystyrene round bottom microwell plates containing LB overnight at 37°C (previously grown overnight in LB and diluted 1,000-fold). Plates were placed in the BioTeK BioSpa 8 automated incubator (Agilent), and OD595 was measured every 30 minutes by the BioTek Citation 3 plate reader (Agilent) for 24h. Growth curves were extracted and area under the curve (AUC) calculated by using an in-house Python code (https://github.com/Cabreiro-Lab/cell_dynamics). Growth curves and stats were performed in Prism 8 (v8.4.0) and in RStudio.

#### Bacterial overexpression mutant generation

We used strains from the ASKA library, based on the *E. coli* K-12 strain^74^. The expression of the ORF of interest is under the control of an IPTG-inducible promoter (isopropyl β-D-1-thiogalactopyranoside) on the plasmid pCA24N carrying chloramphenicol resistance. Clones overexpressing *btuB* and *tonB* were grown in LB broth supplemented with 30 µg/mL of chloramphenicol at 37°C shaking at 200 rpm, plasmids were then extracted with the kit Miniprep GenElute (Simga Aldrich PLN350) and resuspend in water. Plasmids were transformed into strains of interest using the TSS enhanced protocol^72^. Once the transformation was confirmed by PCR, we grown these strains in LB broth supplemented with 1 mmol/L of IPTG at 37°C shaking at 200 rpm for 16 hours.

#### UV-irradiation of bacteria

Bacteria strains were irradiated with UV to inactivate them^75^. To prepare UV-irradiated *E. coli*, an overnight culture was grown in LB broth at 37°C with shaking at 200 rpm for 16 hours. A CL-1000 UV crosslinker equipped with UV-B lamps was sterilized by wiping with 70% ethanol and irradiating the chamber for 5 minutes alternatively. The overnight culture was diluted in fresh sterile LB at a 1:3 ratio and placed in petri dishes. Plates were placed inside the UV chamber without lids and irradiated for a total of 60 minutes, swirling every 10 minutes to ensure uniform exposure. To prevent heat shock-induced bacterial death, the chamber was allowed to cool for 5 minutes between intervals. Following UV treatment, bacteria were collected into a new sterile 50 mL Falcon tube, centrifuged at 4000 rpm for 10 minutes at 4°C, and the supernatant was carefully removed. The bacterial pellet was resuspended in LB and placed on NGM plates for worms.

### Protein identification and quantification by LC-MS/MS

#### Bacterial samples preparation

*E. coli* BW25113, wild type, Δ*lon*::*kan*, Δ*htpG*::*kan*, Δ*dnaK*::*kan*, Δ*clpX*::*kan*, Δ*cbpM*::*kan*, Δ*dnaK*Δ*clpX*::*kan* were grown in LB broth overnight at 37°C shaking 200 rpm. NGM plates were seeded with 120 μL of overnight bacterial cultures and lawns were left to grow at 25°C for 2 days. 5 biological replicates were included per condition. Bacteria were collected from plates with PBS 1X buffer using a sterile glass scraper in Diagenode tubes. Samples were centrifuged at 14000 rpm for 90s at room temperature. The supernatant was removed, and pellets were resuspended with lysis Buffer (8 mol/L urea, 20 mmol/L hepes pH 8). Samples were flash frozen in liquid nitrogen and kept on ice from this point onward. Pellets were then lysed via sonication for 5 minutes at 100% amplitude by using the sonicator waterbath QSonica Q700. Samples were centrifuged at 20000g for 15 minutes at 4°C to separate the cellular debris and proteins. Supernatants containing the extracted protein were transferred to clean tubes and protein concentrations were determined by the Quick start Bradford protein assay (Biorad) at 565 nm. The BSA was used for standard curves. We proceeded to two proteomic analyses, the first one with *E. coli* BW25113, wild type, Δ*lon*::*kan*, Δ*htpG*::*kan*, Δ*dnaK*::*kan*, Δ*clpX*::*kan*, Δ*cbpM*::*kan.* The second one has been proceeded with *E. coli* BW25113 wild type, Δ*dnaK*::*kan*, Δ*clpX*::*kan*, and Δ*dnaK*Δ*clpX*::*kan*.

#### Worm samples preparation

N2 worms were cultivated on NGM plates seeded with *E. coli* BW25113 wild type for 5 days. Eggs were harvested and L1 were seeded on NGM seeded with bacterial strains of interest that have been incubated 2 days at 25°C. 5 biological replicates were included per condition. After 4 days, worms were harvested and washed 5 times with PBS 1X buffer and transferred in Diagenode tubes. Worms were then resuspended in the lysis buffer (8 mol/L urea, 20 mmol/L hepes pH 8.0). Samples were flash frozen with liquid nitrogen then sonicated 2 times 5 minutes at 100% amplitude by using the sonicator waterbath QSonica Q700. Samples were centrifuged 20 min 20000 rpm 4°C. Supernatants containing the extracted protein were transferred to clean tubes and protein concentrations were determined by the Quick start Bradford protein assay (Biorad) at 565 nm. BSA was used for standard curves.

#### Sample preparation for bacterial proteomics

Protein samples (100 µg per sample) were processed using an in-solution digestion procedure. Briefly, samples were sequentially reduced and alkylated at room temperature and in the dark, to final concentrations of 10 mmol/L dithiothreitol (DTT) and 50 mmol/L 2-chloroacetamide (2-CAM), respectively. Samples were diluted two-fold for the first analysis with 20 mmol/L HEPES (pH 8.0), reducing the urea concentration to 4 mol/L, and diluted 8-fold for the second analysis, reducing the urea concentration to 1 mol/L. This was followed by the addition of 2 µg of trypsin (Promega, V528A) and incubation overnight for the first analysis. For the second analysis, an initial LysC (Wako, 121-05063) digestion at a 1: 500 proteases to protein ratio, for 5 hours at 37°C. Samples were then further diluted to a final urea concentration of 2 mol/L with 20 mmol/L HEPES (pH 8.0), followed by the addition of trypsin (Serva, 37286.03) at 1:50 protease to protein ratio. Samples were incubated at 37°C for 16 hours. The digestion of the first analysis was stopped by acidification with a final concentration of 1% trifluoroacetic acid (TFA) against 0,2% for the second one and protein digests were desalted using Glygen C18 spin tips (Glygen Corp, TT2C18.96). Tryptic peptides were eluted with 60% acetonitrile, 0.1% formic acid (FA). Eluents were dried by vacuum centrifugation.

#### Sample preparation for worm proteomics

Protein samples (100µg/sample in 8M urea) were processed using an in-solution digestion procedure. Briefly, samples were sequentially reduced and alkylated at room temperature and in the dark, to final concentrations of 10mM dithiothreitol (DTT) and 50mM 2-chloroacetamide (2-CAM), respectively. Samples were diluted 8-fold with 20mM HEPES (pH 8.0), reducing the urea concentration to 1.5M. This was followed by addition of 2µg of trypsin (Promega, V528A). Samples were incubated over-night at 37°C. The digestion was stopped by acidification with 10% trifluoroacetic acid (TFA) to a final concentration of 1% and protein digests were desalted using Glygen C18 spin tips (Glygen Corp, TT2C18.96). Tryptic peptides were eluted with 60% acetonitrile, 0.1% formic acid (FA). Eluents and dried by vacuum centrifugation.

#### Liquid chromatography-tandem mass spectrometry (LC-MS/MS) analysis

Dried tryptic digests were re-dissolved in 0.1% TFA and each sample injected at 2μg LC-MS/MS analysis was performed using an Ultimate 3000 RSLC nano liquid chromatography system (Thermo Scientific) coupled to a coupled to a Q-Exactive mass spectrometer (Thermo Scientific) via an EASY spray source (Thermo Scientific). For LC-MS/MS analysis re-dissolved protein digests were injected and loaded onto a trap column (Acclaim PepMap 100 C18, 100 μm × 2cm) for desalting and concentration at 8 μL/min in 2% acetonitrile, 0.1% TFA. Peptides were separated on-line to an analytical column (Acclaim Pepmap RSLC C18, 75 μm × 75 cm for the bacterial samples, and C18, 75 μm × 50 cm for the worm samples) at a flow rate of 200 nL/min and 250 nL/min for the bacteria and worm samples respectively). For bacteria samples, peptides were separated using a 120 minutes gradient, 4-25% of buffer B for 90 minutes followed by 25-45% buffer B for another 30 minutes (composition of buffer B – 80% acetonitrile, 0.1% FA). For worm samples, peptides were separated using a 90 minutes gradient, 1-22% of buffer B for 60 minutes followed by 22-44% buffer B for another 30 minutes (composition of buffer B – 75% acetonitrile, 5% DMSO and 0.1% FA). Eluted peptides were analyzed by the mass spectrometer operating in positive polarity using a data-dependent acquisition mode. Ions for fragmentation were determined from an initial MS1 survey scan at 70000 resolution for bacterial samples and 120000 for worm samples, followed by HCD (Higher Energy Collision Induced Dissociation) of the top 12 most abundant ions for bacteria samples and 30 most abundant ions for worm samples at 17500 resolution. MS1 and MS2 scan AGC targets were set to 3e6 and 5e4 for maximum injection times of 50ms and 50ms respectively. A survey scan m/z range of 375 – 1800 was used, normalized collision energy set to 27%, charge exclusion enabled with unassigned and +1 charge states rejected and a minimal AGC target of 1e3. Dynamic exclusion was set to 45-50 seconds.

#### Data analysis for proteomics

Raw mass spectrometry files were processed in MaxQuant using *E. coli* K12 (version 20200915) or *C. elegans* (version 20210628) databases and label-free quantification with minimum LFQ ratio of 1. LFQ values were then inputted in Perseus for statistical analysis (version 1.6.2.3 for the bacterial samples analysis and version 1.6.10.43 for the worm samples) to obtain statistical and bioinformatic analysis, as well as for visualization. The data matrix was filtered based on categorical columns to remove reverse decoy hits, potential contaminants and protein groups which were ‘only identified by site’. Gene annotations were done by using *E. coli* K12 (version 20200915) or *C. elegans* (version 20210628) GOBP, GOMF, GOCC, and KEGG database. Data were log2 transformed. The 5 biological replicates for each mutant were then pooled, compared to each other and visualized as Volcano plots. Volcano plots were generated based on LFQ intensities with the following settings: T-test; side: both; number of randomizations: 250; preserve grouping in randomizations: <none>; FDR: 0.05; S0: 0.1. Then, significant differences between mutants were exported for a Hierarchical clustering analysis (HCA). This was carried out after filtering rows based on a minimum of two valid values in at least one group, Z-scoring of values in rows. The HCA was generated with the following settings for both rows tree and columns tree: distance: Euclidean; linkage: average; constraint: none; preprocess with k-means selected (number of clusters: 300; maximal number of iterations: 10; number of restarts: 1). Further data representation and plotting was carried out in R programming language.

Given that both Δ*clpX* and Δ*dnaK*Δ*clpX* behave in a similar way opposed to the Δ*dnaK* deficient strain, we subtracted the differences between groups to study the proteins that were unique to each cluster. We were specifically interested in the set of proteins that were downregulated in Δ*dnaK* opposed to the upregulated in Δ*dnaK*Δ*clpX*, we used the double mutant as a control and subtracted the proteins found in Δ*dnaK*. Therefore, the effects shown in the distinct proteins between both groups can be described as the unique signature of the differential proteostasis capabilities of both groups. In a similar way, the set of proteins expressed in Δ*dnaK* but not in the other groups was studied using the same logic. Thus, the set of proteins that conferred protein stability was also captured here. Groups were extracted from the significant proteins using R programming language and the UpSet library v. 1.4.0.

#### Western blot

Worms were grown on plates seeded with *E. coli* BW25113, Δ*lon*::*kan*, Δ*htpG*::*kan*, Δ*dnaK*::*kan*, Δ*clpX*::*kan*, Δ*cbpM*::*kan*, Δ*hscA*::*kan*, Δ*dnaJ*::*kan*, Δ*hybE*::*kan* from the L1 to day1 adult stage at 20°C. 75 worms were collected in 100 μL 1X SDS loading buffer. Then samples were boiled 5 minutes at 95°C at 1400 rpm, sonicated for 5 minutes at 100% of amplitude by using the sonicator waterbath QSonica Q700, and boiled again for 5 minutes at 95°C at 1400 rpm. Samples were then centrifuged for 5 minutes at 14000 rpm. For the western blot, proteins from the lysate worms were separated by size using an Invitrogen precast SDS-Page gel 4-12%. Separated proteins were transferred on a nitrocellulose membrane by a dry blotting system (iBlot 2 dry blotting system) with a setting according to manufacturer’s instructions. For the detection of GFP, mCherry and Tubulin, the membranes were probed with primary Mouse monoclonal antibodies anti-GFP at a 1:5000 dilution (clone JL-8), anti-mCherry at a 1:2000 dilution (clone 1C51), anti-alpha tubulin at a 1:10000 dilution (clone B-5-1-2) respectively. Then membranes were exposed to the secondary antibody, Li-Cor anti-Mouse 800CW/680 from Donkey at a 10000 dilution.

The intensity of each GFP band was normalized by the intensity of its corresponding mCherry and Tubulin bands. 3-4 biological replicates were included per condition. Statistical analysis was done by using a one-way ANOVA with multiple comparisons (Tukey’s multiple comparison test) with the software PRISM8 (version 8.4.0).

#### Nematode fluorescence microscopy

PP607 worms (UBV worms) were cultivated on NGM plates seeded with *E. coli* BW25113 wild type for 5 days. Eggs were harvested and L1 were seeded on NGM previously seeded with bacterial strains of interest incubated 2 days at 25°C. After 4 days at 20°C, a minimum of 11 worms were anesthetized with 2% levamisole on NGM plates and were imaged under a 40x objective using a Zeiss Axio Zoom V16 microscope system equipped with an AxioCam MRm camera operated by Zen 2 software (Zeiss). Either the GFP filterset (excitation: 450-490 nm; emission: 500-550 nm) or the mCherry filterset (excitation: 559-585 nm; emission: 600-690 nm) was used. All images were exported in CZI format and fluorescence levels were quantified using Volocity 5.2 software (PerkinElmer) run on a Surface tablet (Microsoft). The fluorescence intensity of worms was calculated as the pixel density of the entire cross-sectional area occupied by worms from which the pixel density of the background had been subtracted. 3 independent replicates were carried out with a minimum of 11 worms imaged per condition per replicate. The fluorescence intensity was calculated automatically by setting a minimum threshold intensity that excluded the background.

3-6 biological replicates were included per condition. Statistical analysis was done by using a one-way ANOVA with multiple comparisons (Tukey’s multiple comparison test) with the software GraphPad PRISM8 (version 8.4.0).

## DATA AVAILABILITY

The raw sequences for the transcription profiles of the mono-association experiments with *C. elegans* and the *E. coli* pangenome reported in this study can be accessed in GSE315953. The raw proteomics profiles reported in the experimental validation can be accessed in PRIDE under the IDs PXD071769, PXD071818 and PXD071867. The data and a stable version of the code comprising the pangenome analysis, protein embeddings, strain embeddings and the processed transcriptome data can be accessed in Zenodo under the DOI 10.5281/zenodo.18221759.

## ACKNOWLEDGEMENTS

F.C. was supported by the Wellcome Trust/Royal Society (102532/Z/12/Z and 102531/Z/13/A), the DFG German Research Foundation (EXC 2030 -390661388) and (IPFP-B02 Filipe Cabreiro) of the Center for Molecular Medicine Cologne. D.M.M was supported by the Leverhulme Trust (RPG-2022-299). HMD was supported by i-HEAD“ (PB-NRW-2022-ANTRAG-025). We acknowledge computational resources and support provided by the Imperial College Research Computing Service (http://doi.org/10.14469/hpc/2232). We would like to thank Saul Moore for helping with developing a method to select focused microscope images, Jennifer van der Laan and Evgeny Galimov for technical support and Thorsten Hoppe and Franziska Ottens for UPS-related scientific discussions. We would like to express our gratitude to our colleagues and to Dr. Mária Džunková which helped to shape this manuscript.

## AUTHOR CONTRIBUTIONS

D.M.M., F.C. conceptualized the research. D.M.M. performed the computational analysis of the pangenome. D.M.M, A.A., C.B., H.M.D., J.W. and F.C. analyzed the data. A.A. performed the reporter validation and transcriptome studies. C.B., A.Z., J.W. and F.C performed the experiments for the worm proteostasis. A.I. and L.G. sequenced transcriptomes and genomes. I.K., G.R, A.M. and H.K. performed proteomics identification and analysis. D.M.M., and F.C. wrote the manuscript. D.M.M., C.B., A.A. and F.C. participated in editing the manuscript. D.M.M. and F.C. participated in the interpretation of the main findings. D.M.M and F.C. supervised the research. All authors read and approved the final manuscript.

## SUPPLEMENTAL FIGURE LEGENDS

**Figure S1.**
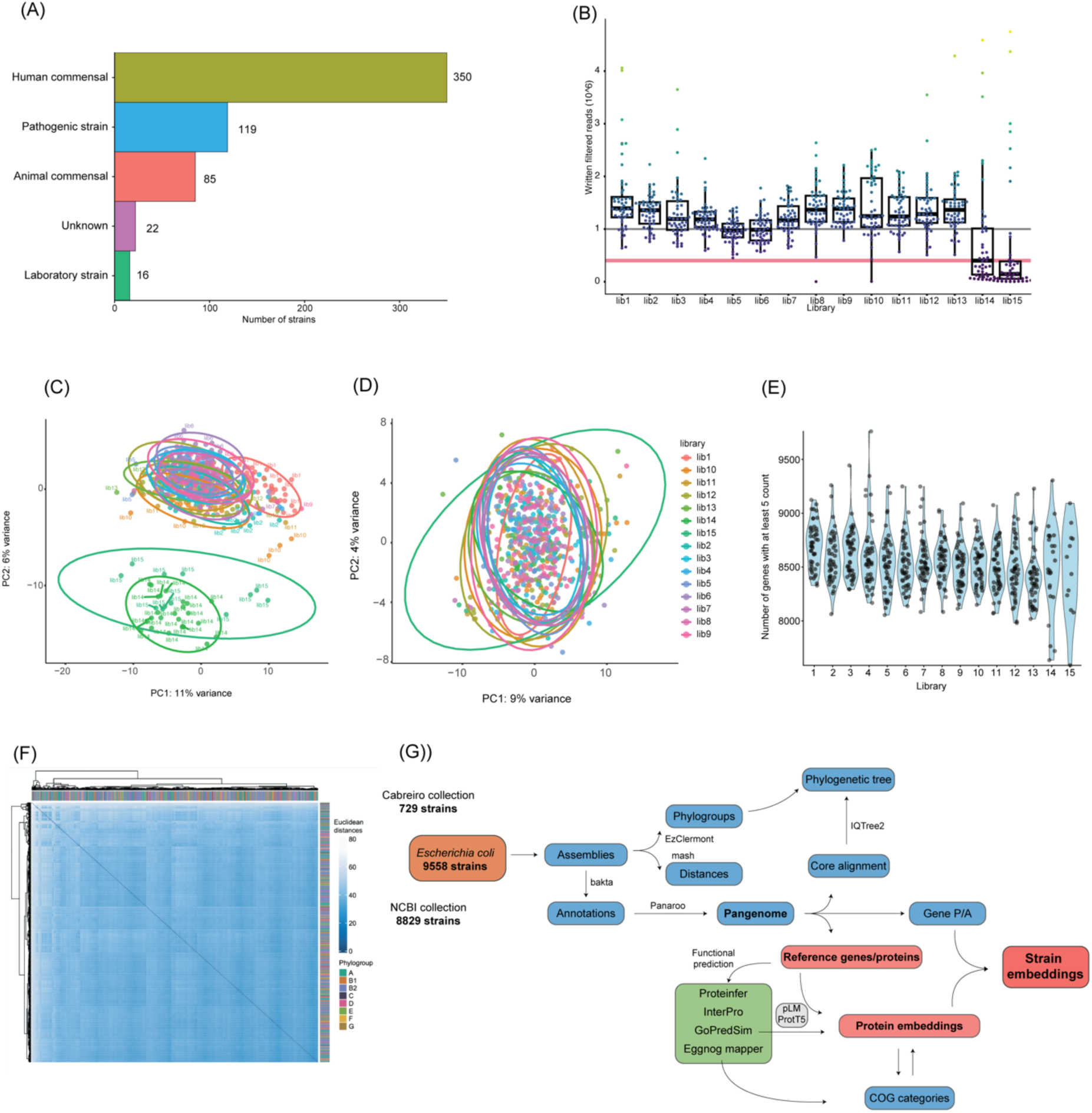
Strain composition and RNA-seq quality control. (A), Number of *E. coli* strains in the panel classified as human commensal, human pathogenic, animal commensal, laboratory strains or of unknown origin. (B), Read-count distribution per library for the high-throughput RNA-seq experiment, generated on a NextSeq 2000 P3 run (1.2 × 10⁹ total reads). (C), PCA of *C. elegans* transcriptomes after outlier removal, showing batch-driven separation with samples from libraries 1, 14 and 15 forming distinct clusters. (D), PCA of the same dataset after batch correction, showing no library-driven separation. (E), Violin plot of the number of genes detected per worm with at least 5 normalized counts. Each dot represents a worm sample; data is represented by sequencing library. (F), Heat map of pairwise Euclidean distances based on DESeq2-normalized *C. elegans* gene expression, with hierarchical clustering of libraries with dark blue-to-light blue showing close-to-distant relationships between samples. PCA data and Euclidean distances are showing the VST normalized data from the transcriptional profiles.

**Figure S2.**
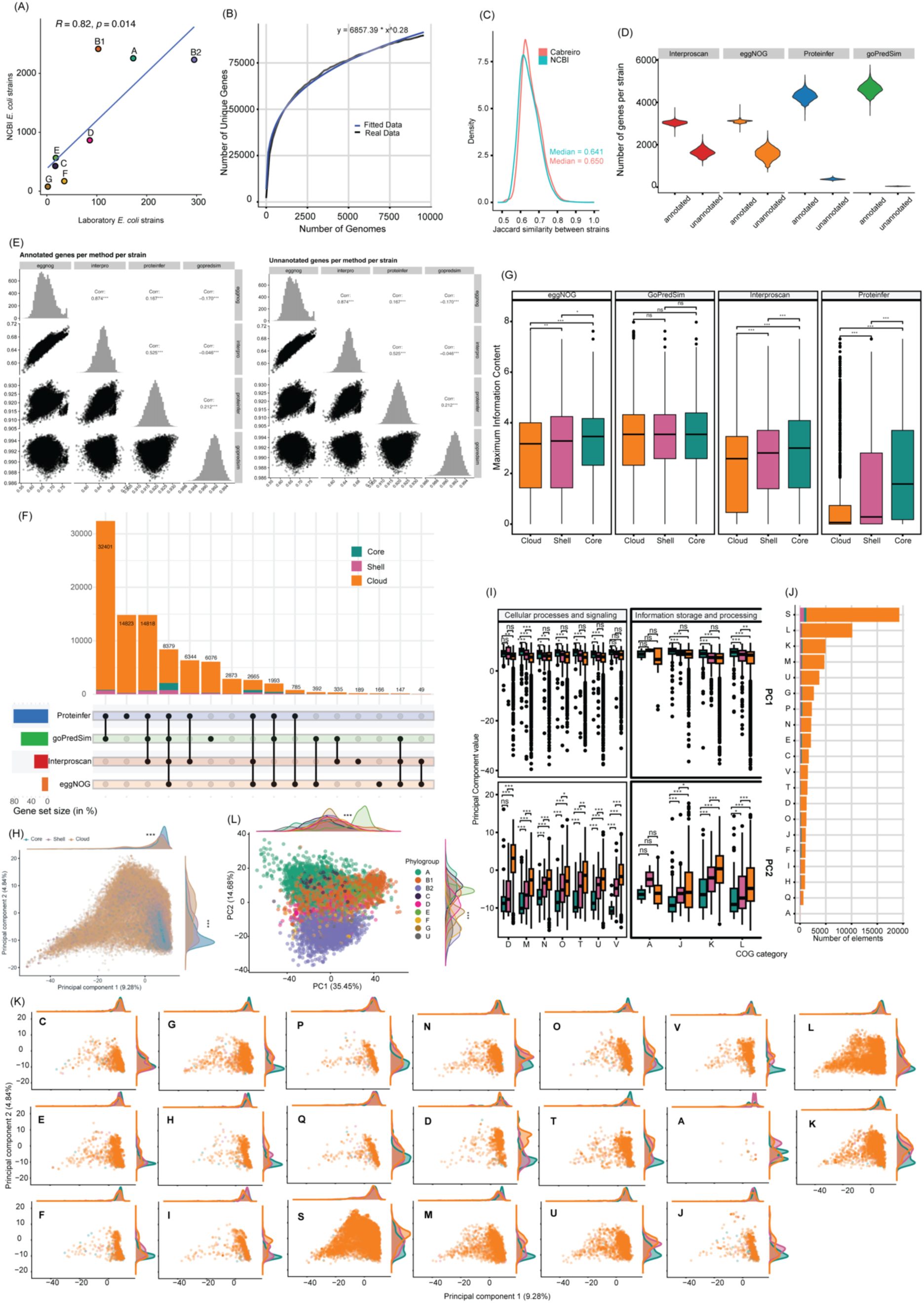
Functional annotation performance and embedding structure of the *E. coli* pangenome. (A), Phylogroup frequency Pearson correlation between the laboratory *E. coli* trains and an NCBI *E. coli* collection (R = 0.82, P = 0.014). (B), Pangenome accumulation curve for the 9,558 *E. coli* genomes. The x-axis shows the number of genomes progressively added to the analysis, and the y-axis shows the cumulative number of unique genes observed with increasing pangenome size. (C), Distribution of the pairwise Jaccard similarity between *E. coli* strains from NCBI and the Cabreiro lab collection. Median values per collection are represented. (D), Violin plots representing the fraction of genes annotated by different functional annotation tools: Interproscan, eggNOG-mapper, Proteinfer and GOPredSim. (E), Pearson correlation of annotated (left) and unannotated (right) fraction per genome and per method (n=92,435). (F), UpSet diagram of the number of genes with a GO term annotation by each one of the functional annotation methods. Dots in the lower part describe which tool or tools are being considered in each case. Genome partition is represented as a color in the bar-plot. (G), Maximum information content for the GO terms annotated by each method per genome partition (n=1,347-60,152). (H), PCA projection of the protein embeddings from the linear reference from the *E. coli* pangenome. Colors represent the genome fractions of core, shell and cloud (n=92,244). (I), Principal component values of gene embeddings stratified by COG functional category and pangenome class (n=2-9,802). (J), Distribution of genes across COG categories, partitioned into core, shell and cloud components. Bars show, for each COG category, the number of genes assigned to the core, shell and cloud genome. (K), PCA projections of protein embeddings split per genes classified in the different COG categories. (L), PCA projections of the strain embeddings colored by the main *E. coli* phylogroups (n=9,558). Data shown in (G), (I), (H) and (L) were tested with two-tail pairwise T-test and significance is expressed as *P < 0.05, **P < 0.01, ***P < 0.001, NS P > 0.05.

**Figure S3.**
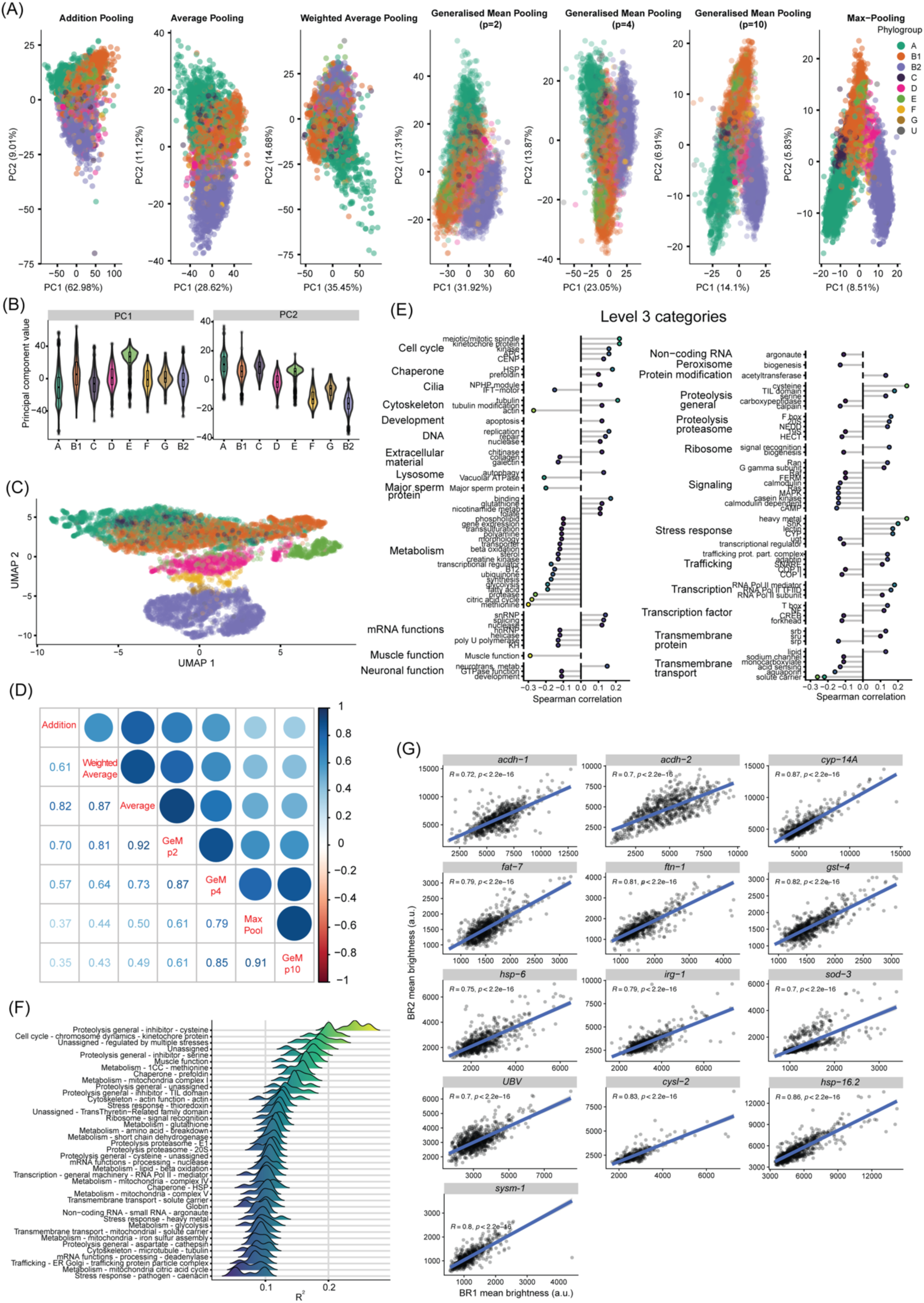
Construction and structure of *E. coli* strain embeddings. (A), PCA of strain embeddings for the 9,558 *E. coli* strains, built using seven pooling strategies: addition of gene embeddings, average of gene embeddings, weighted average of gene embeddings, generalized mean pooling with parameter p set to 2, 4 and 10, and max-pooling (see Methods for a detailed description). Points are colored by phylogroup. (B), Violin plots showing the distributions of PC1 (left) and PC2 (right) coordinates of strain embeddings, calculated as the average of gene embeddings, across phylogroups. (C), UMAP projection of strain embeddings for all 9,558 *E. coli* strains, revealing distinct clusters colored by phylogroup. (D), pairwise correlation of strain pooling strategies calculated from Procrustes test, every single correlation is significant. (E), Bubble plot of the significant Spearman correlation coefficients (ρ) between PC2 of the *E. coli* strain embeddings and WormCat functional scores across all significant worm functional level 3 categories. Worm categories are grouped by level 1 hierarchy. (G) Pearson correlation representation of two biological replicates per gene transcriptional reporter in *C. elegans* (n=589). (F), Mean Pearson correlation (R^2^) from the Random Forest cross-validated prediction of worm phenotype using the seven pooling strategies.

**Figure S4.**
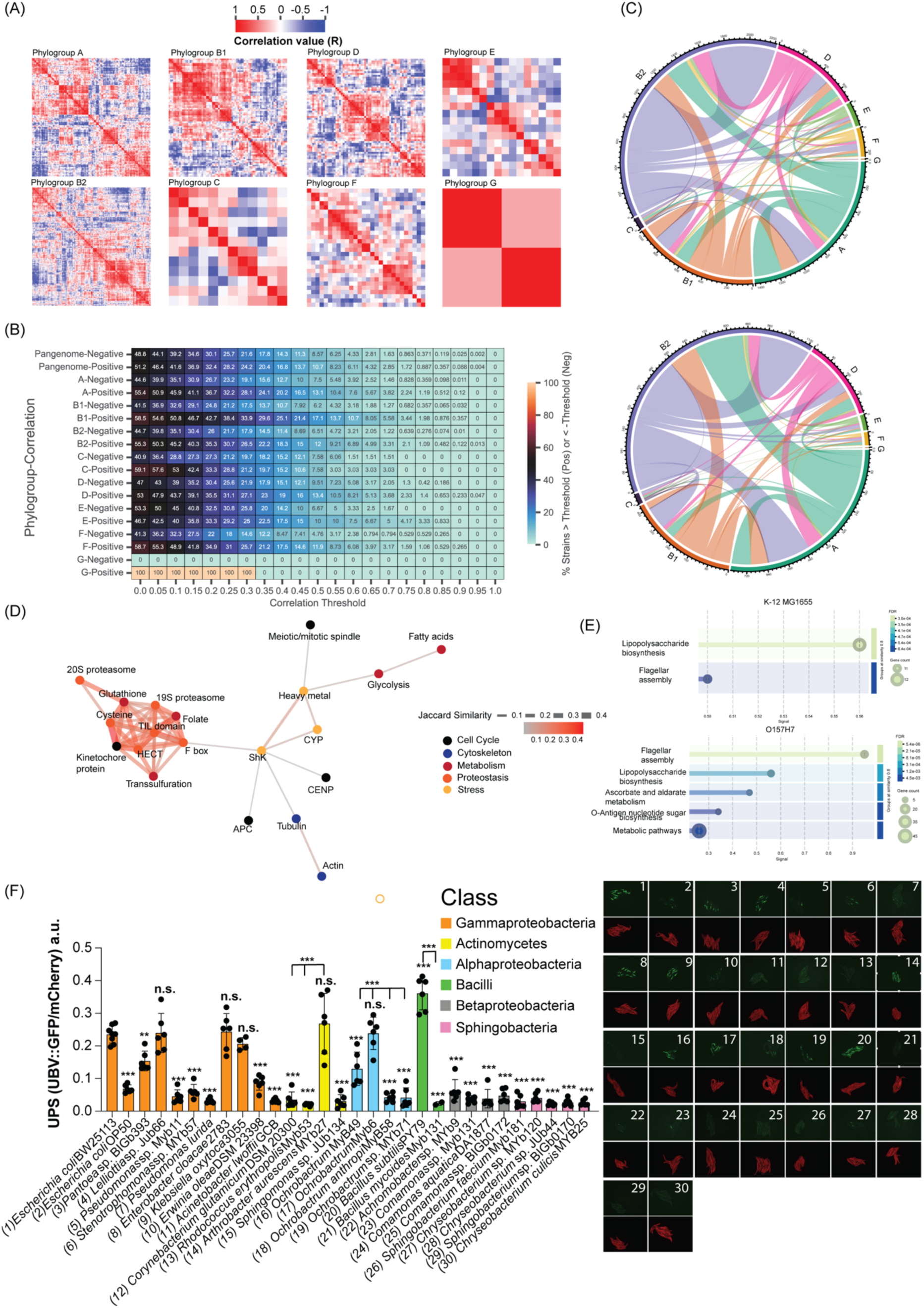
Associations between strain embeddings, WormCat functional responses and gene reporter phenotypes. (A), Pairwise Pearson correlation coefficients between *E. coli* strains separated by phylogroup (n=589). (B), Heat map representing the percentage of positive and negative correlations within *E. coli* phylogroups given a range of correlation thresholds (x-axis) (n=589). (C), Chord plots representing the within and between strain correlations between the main *E. coli* phylogroups for the positive (left) and positive (right) correlations (n=8-13). (D), Worm physiological processes driven by shared genetic drivers from *E. coli* measured by their permutational Jaccard distances. The network represents co-occurrence of the processes, with nodes representing worm pathways and edges representing the Jaccard similarity. (E), KEGG functional enrichment analysis from the shared genes affecting the proteostasis-proteolysis-1CC metabolism cluster from (D) using *E. coli* K-12 (top) or O157H strains (bottom) as backgrounds in STRING DB. (F) Left, normalized UPS fluorescence quantification of worms fed individually on several bacterial species and strains, where color represents each bacterial phylum (n=2-8). Right, representative fluorescence images of the Ub(G76V)–GFP and constitutively expressed mCherry reporters. Correspondence between the quantification and representative images is done by a numeric code. *P < 0.05, **P < 0.01, ***P < 0.001, NS P > 0.05, one-way ANOVA.

**Figure S5.**
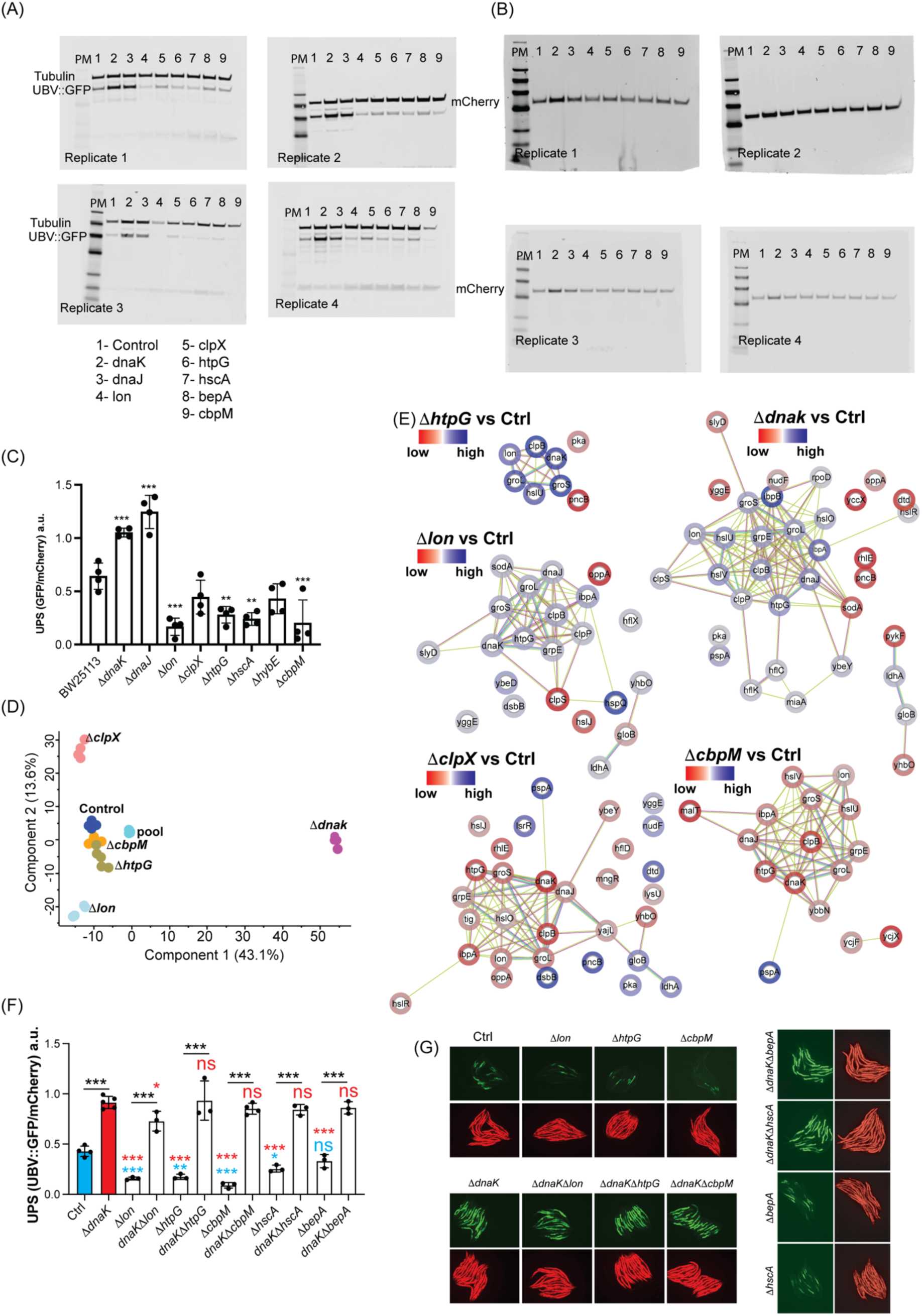
Proteostasis at the host level is regulated by bacterial chaperones. (A-B), Western blot analysis of Tubulin, UBV::GFP (A) and mCherry (B) expression in from worms grown in *E. coli* chaperone and protease mutants. Each replicate (1-4) shows worm protein expression in various *E. coli* mutants: 1-BW25113 (control), 2-Δ*dnaK*, 3-Δ*dnaJ*, 4-Δ*lon,* 5-Δ*clpX,* 6-Δ*htpG*, 7-Δ*hscA*, 8-Δ*bepA*, and 9-Δ*cbpM*. Tubulin serves as a loading control. (C), Ratio of the quantification of UBV::GFP protein expression over mCherry protein expression (n=4) (*P < 0.05, **P < 0.01, ***P < 0.001, NS P > 0.05, one-way ANOVA). (D) Principal Component Analysis of the protein expression profile from proteomics analysis of *E. coli* BW25113 control and Δ*dnaK*, Δ*clpX*, Δ*lon*, Δ*htpG* and Δ*cbpM* mutants. (E), Network representation from differentially expressed proteins from for the comparisons to control BW25113 to Δ*dnaK*, Δ*clpX*, Δ*lon*, Δ*htpG* and Δ*cbpM* mutants; protein-protein interactions extracted from STRING database. Colours represent high (blue) and low (red) expression compared to the control strain BW25113. (F), Normalized UPS fluorescence quantification of worms fed *E. coli* BW25113 and indicated bacterial chaperone and protease mutants. Stats are represented as coloured stars, black for the double Δ*dnak* mutant vs Δ*dnaK* single mutant, blue for the comparison against the control strain BW25113, and red for the comparison against Δ*dnaK* mutant. (n=3-6, *P < 0.05, **P < 0.01, ***P < 0.001, NS P > 0.05, One-way ANOVA). (G), Representative fluorescence images of the Ub(G76V)–GFP and constitutively expressed mCherry reporters from worms fed on each bacterial mutant tested in (D) and (F).

**Figure S6.**
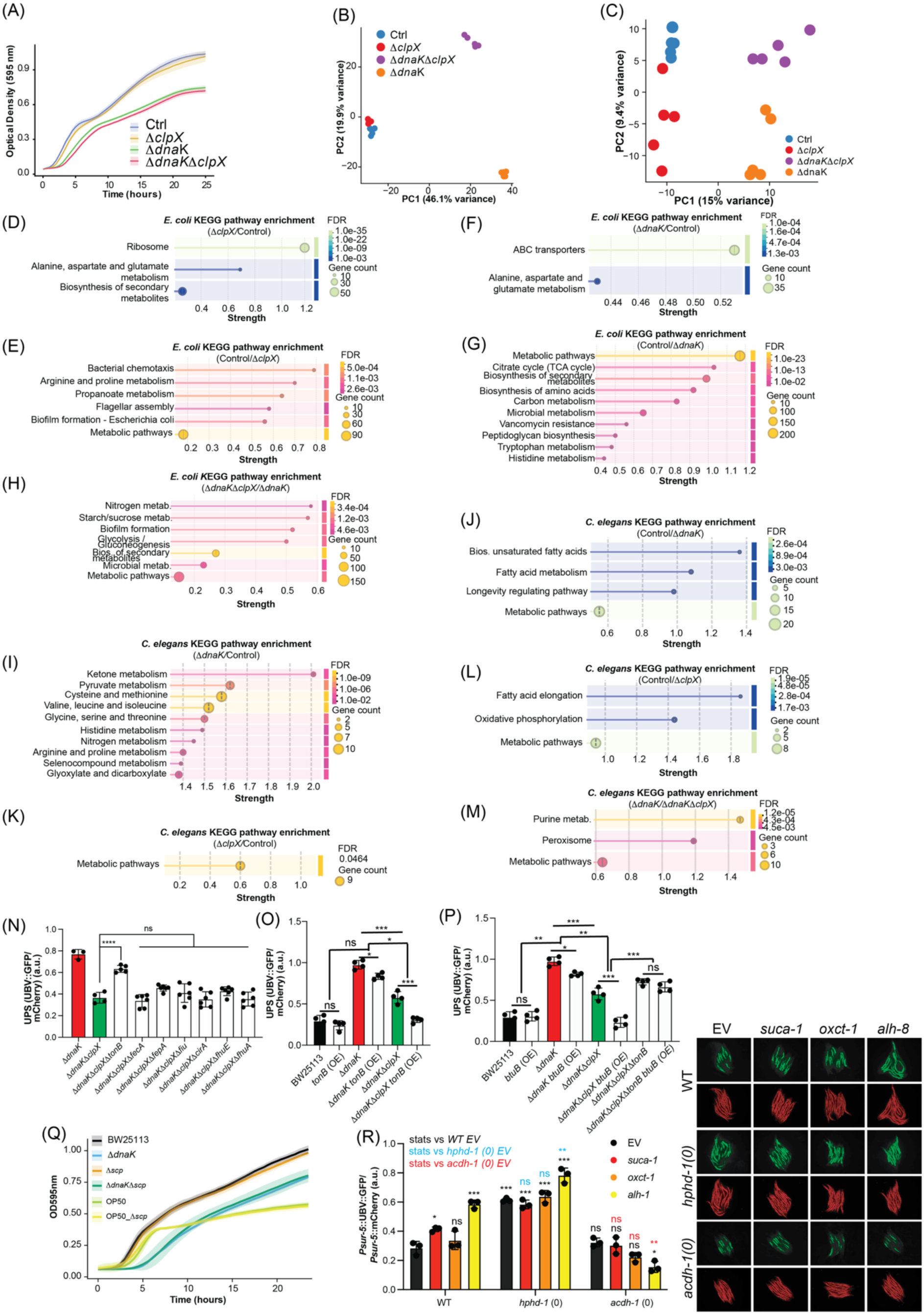
Bacterial chaperones and proteases drive proteostasis regulation in the host through propionate and V12 metabolism. (A), Growth curves showing optical density at 595 nm plotted over time (hours) of wild-type BW25113 (control), Δ*clpX*, Δ*dnaK* and Δ*dnaK*Δ*clpX* double-mutant *E. coli* strains. (B), Prinicipal Component Analysis of the protein expression from proteomics data obtained from *E. coli* strains BW25113 control and Δ*clpX*, Δ*dnaK* and Δ*dnaK*Δ*clpX* mutant strains. (I) Principal Component Analysis of the protein profiles from proteomics data from *C. elegans* fed *E. coli* BW25113 (Control) and mutants Δ*dnaK*, Δ*clpX* and Δ*dnaK*Δ*clpX*. (D-M) KEGG pathway enrichment analysis of proteomics data for the comparisons shown in *E. coli* mutants or worms fed the respective *E. coli* mutants using STRING DB. (N), Normalized UPS fluorescence quantification of worms fed control BW25113 and indicated *E. coli* knockout mutant strains (n=3-5).(O-P), Normalized UPS fluorescence quantification of worms fed control BW25113 and *E. coli* knockout mutant strains or overexpressing the indicated genes (n=4). (Q), Growth curves showing optical density at 595 nm plotted over time (hours) of wild-type *E. coli* BW25113 (control), Δ*dnaK*, Δ*scp*, Δ*dnaK*Δ*scp* double-mutant. *E. coli* OP50 and the OP50 mutant Δ*scp* correspond to the green and yellow lines. (R), Left, Fluorescence quantification of the reporters UBV::GFP and mCherry ratio in the worm strains from right. Right, fluorescence images of the UBV::GFP and mCherry reporters from *C. elegans* N2 strain (Control), *acdh-1*(0), *hphd-1*(0) in combination with KO Empty Vector (EV), *suca-1*, *oxct-1*, *alh-8* strains fed on *E. coli* BW25113. For enrichment plots, color represents the FDR values and circle size the number of genes per category. Statistics are represented as *P < 0.05, **P < 0.01, ***P < 0.001, NS P > 0.05, using one-way ANOVA for pairwise interactions and two-way ANOVA for nested conditions.

## SUPPLEMENTARY INFORMATION

### Supplemental Data

Table S1: Metadata for the E. coli strains used for training and in the laboratory Table S2: RNA-seq normalized counts into level 3 categories from WormCat

Table S3: Pangenome summary statistics of gene families found in the 9,558 E. coli strains Table S4: PCA coordinates for average strain embeddings

Table S5: Spearman correlation analysis for PC2-level3 pairings Table S6: Worm reporter brightness data, normalized

Table S7: Network topology of protein-protein interactions in E. coli with effect size Table S8: Worm brightness screen for E. coli genes involved in proteostasis

Table S9: Worm brightness data for experiments in Figures 5 and 6 Table S10: Bi-partite graph data from the proteomics experiment

Table S11: List of worm reporters used for the screening with the pathway readout. Table S12: List of bacterial mutants and primers used to test co-proteostasis

